# Oxidized phosphatidylinositol impairs lysosomal membrane repair to promote ferroptosis

**DOI:** 10.64898/2026.09.10.750804

**Authors:** Shota Fukuyama, Ping-Yen Hsieh, Yuka Jinnouchi, Yuta Matsuoka, Eisho Kozakura, Tomohiro Segawa, Hiroaki Matoba, Hideto Naka, Kyoko Mine, Yuuna Shiraiwa, Miku Yoneda, Takumi Udo, Nao Kato, Hiroki Nakanishi, Eikan Mishima, Yuki Sugiura, Mirinthorn Jutanom, Masami Abe, Pakawit Lerksaipheng, Marcus Conrad, Go Hirai, Ken-ichi Yamada

## Abstract

Ferroptosis is a regulated form of cell death driven by iron-dependent and unrestrained lipid peroxidation, which generates phospholipid hydroperoxides that cause membrane rupture. Oxidized phospholipid species, including oxidized arachidonic acid-containing phosphatidylethanolamines (PE), are abundant during ferroptosis. However, previous studies have examined only a limited number of lipid species, and it remains unclear which oxidized phospholipids consistently arise across distinct cell types and ferroptosis-inducing conditions. Here, we comprehensively profiled oxidized phospholipids generated during ferroptosis across multiple cell lines and animal models. We identified PE 18:0_20:4;O3 and phosphatidylinositol (PI) 18:0_20:4;O3 as oxidized phospholipid species that are consistently detectable across all tested ferroptosis-inducing conditions. Furthermore, oxidized phosphatidylinositol impairs the phosphoinositide-initiated membrane tethering and lipid transport (PITT) pathway, a key mechanism for lysosomal membrane repair. These results indicate that the oxidized phospholipids identified here may serve as markers of ferroptosis while also acting as bioactive mediators that compromise lysosomal membrane homeostasis and repair.

## Introduction

Ferroptosis is a regulated form of cell death characterized by iron-dependent peroxidation of membrane phospholipids and by morphological and biochemical features distinct from those of other cell death modalities, including apoptosis^1–3^. Ferroptosis contributes to the pathogenesis and progression of diverse disorders, including acute kidney injury and neurodegenerative diseases^4–7^, whereas its induction has emerged as a potential therapeutic strategy for treatment-resistant cancers^8, 9^. Therefore, elucidating the molecular mechanisms that govern the initiation, propagation, and suppression of ferroptosis is important both for understanding disease pathogenesis and for developing new therapeutic approaches.

The execution of ferroptosis is driven by the peroxidation of membrane phospholipids containing polyunsaturated fatty acids (PUFAs) and the resulting increase in phospholipid hydroperoxides^10–12^. In particular, oxidized phosphatidylethanolamine (oxPE) species containing arachidonic acid have been identified as prominent ferroptosis-associated lipids^10^. Beyond PE, comprehensive lipidomic analyses have further shown that ferroptosis is accompanied by increases in diverse oxidized phospholipid species across multiple phospholipid classes^13, 14^. However, oxidized phospholipid profiles are dependent on the analytical platform of oxidized lipid detection and may vary among cell types and ferroptosis-inducing conditions, and whether specific molecular species are reproducibly increased across different experimental settings remains unclear. Moreover, it is unknown whether individual oxidized phospholipids merely increase as products of lipid peroxidation (LPO) or actively promote ferroptosis by perturbing intracellular membrane homeostasis.

Recent studies have implicated multiple organelles, including the endoplasmic reticulum^15^, mitochondria^16^, peroxisomes^17^ and lysosomes^18^, in the initiation and propagation of ferroptosis^19^. Lysosomes are particularly important because they serve not only as sites of lipid peroxide increase but also as reservoirs of redox-active iron capable of driving LPO. RSL3-induced LPO has been reported to originate in lysosomes, whereas the sequestration or inactivation of lysosomal iron suppresses both LPO and ferroptotic cell death^18^. We previously demonstrated that the increase of lipid-derived radicals in lysosome induces lysosomal membrane permeabilization (LMP), resulting in the release of lysosomal iron into the cytosol and amplifying LPO throughout the cell during ferroptosis^20^. These findings suggest that, in addition to the increased LPO products in lysosomes, the mechanisms that preserve the integrity of oxidatively damaged lysosomal membranes may be critical determinants of ferroptosis.

Here, we extended our previously established analytical platform for oxidized phosphatidylcholine-derived lipids^21^ to include phosphatidylethanolamine (PE), phosphatidylserine (PS), and phosphatidylinositol (PI), and characterized oxidized phospholipid profiles across multiple cell lines and ferroptosis-inducing conditions. Under the conditions examined, PE 18:0_20:4;O3 and PI 18:0_20:4;O3 were consistently increased during ferroptosis. Because the phosphoinositide-initiated membrane tethering and lipid transport (PITT) pathway^22^, a lysosomal membrane repair mechanism, depends on PI metabolism, we further investigated whether ferroptosis-associated PI oxidation affects PITT-mediated maintenance of lysosomal membrane integrity and ferroptosis progression. Our findings establish a functional link between the molecular profile of ferroptosis-associated oxidized phospholipids and organelle membrane repair, and identify disruption of lysosomal membrane homeostasis as a mechanism that promotes ferroptosis.

## Results

### Comprehensive profiling of oxidized phospholipids identifies ferroptosis-associated oxidized phospholipids

To enable comprehensive profiling of oxidized phospholipids, we extended our previously established multiple-reaction monitoring (MRM) library of oxidized phosphatidylcholines (oxPCs) ^21^ to include oxidized phosphatidylethanolamines (oxPEs), phosphatidylserines (oxPSs), and phosphatidylinositols (oxPIs). This extension generated a library of 1,860 oxidized phospholipid species, covering the major phospholipid classes (Fig. 1a).

**Fig. 1.**
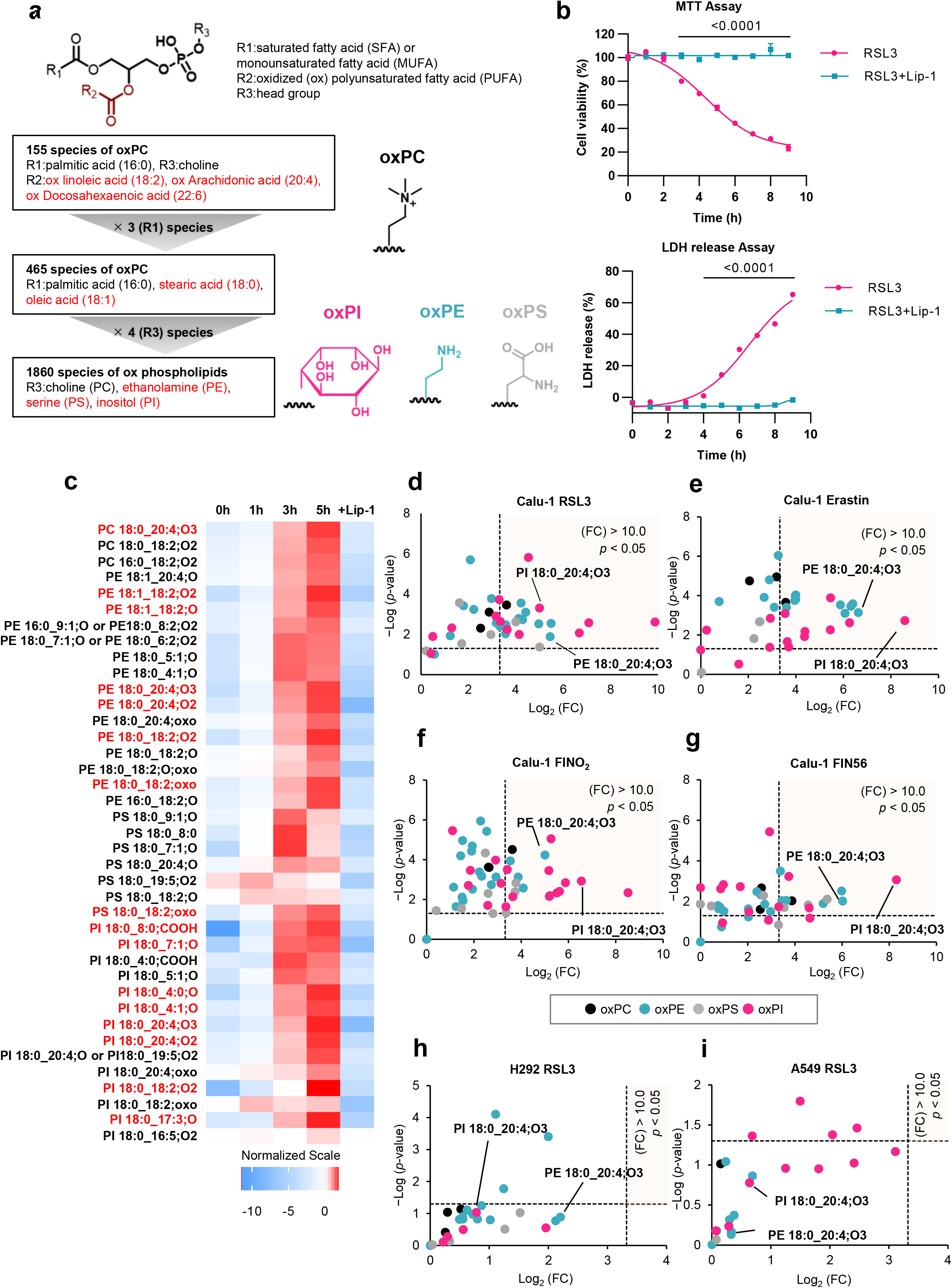
Comprehensive profiling of oxidized phospholipids identifies ferroptosis-associated oxidized phospholipids. **a,** Selection workflow for the comprehensive analysis of oxidized phospholipids generated during ferroptosis. **b,** Viability of Calu-1 cells treated with RSL3 (0.1 µM) ± Lip-1 (1 µM) for the indicated durations (1–9 h). **c,** Fold changes in 39 oxidized phospholipids in Calu-1 cells treated with RSL3 (0.1 µM) for the indicated durations or with RSL3 (0.1 µM) and Lip-1 (1 µM) for 5 h. Among the 22 species that increased significantly by more than tenfold after 5 h of RSL3 treatment, the 16 species whose MS peak areas increased over time are shown in red. The color scale indicates log2-transformed fold changes. **d–g,** Fold changes in oxidized phospholipids in Calu-1 cells treated with RSL3 (0.1 µM) for 5 h (**d**), erastin (10 µM) for 7 h (**e**), FINO2 (10 µM) for 6 h (**f**) or FIN56 (5 µM) for 5 h (**g**). **h,i,** Fold changes in oxidized phospholipids in H292 (**h**) and A549 (**i**) cells treated with RSL3 (0.1 µM) for 5 h. Fold changes in (**c–i)** were calculated from MS peak areas. Data in (**b)** are presented as the mean ± s.d. of three independent experiments. Statistical significance was assessed using Sidak’s multiple-comparisons test. Data in (**c–i)** are presented as log2-transformed mean fold changes from three independent experiments. Statistical significance was assessed using a two-sided unpaired t-test.

We next used this extended library to identify oxidized phospholipid species generated during ferroptosis. Treatment of the human lung cancer cell line Calu-1 with the glutathione peroxidase 4 (GPX4) inhibitor (1*S*,3*R*)-RSL3 (RSL3) reduced cell viability after 3 h, as determined by the MTT assay, and increased LDH release after 4 h (Fig. 1b). Increased C11-BODIPY fluorescence was also detected at 5 h after RSL3 treatment (Supplementary Fig. 1a). All these changes were suppressed by co-treatment with the ferroptosis inhibitor liproxstatin-1 (Lip-1). Based on these findings, treatment of Calu-1 cells with 0.1 μM RSL3 for 5 h was used to induce ferroptosis in subsequent experiments.

LC–MS/MS analysis targeting all 1,860 oxidized phospholipid species detected increased MS peaks corresponding to 687 species in RSL3-treated Calu-1 cells (Supplementary Fig. 1b).

Representative extracted-ion chromatograms (EICs) for PI 18:0_20:4;O3 (*m/z* 933.5335) are shown in Supplementary Fig. 1c. To identify the detected oxidized phospholipids, targeted LC–MS/MS analysis was performed to detect one product ion derived from the PI polar headgroup (*m/z* 241.00) and two product ions derived from the fatty acyl chains (*m/z* 283.25 and 351.20). Detection of all three product ions at the same retention time allowed assignment of the corresponding oxidized phospholipid molecular species (Supplementary Fig. 1d).

By applying a similar analytical procedure to all 687 candidate molecules, we identified 39 oxidized phospholipid species that increased in RSL3-treated Calu-1 cells (Table 1 and Fig. 1c). The MS peak areas for these species were calculated from EICs corresponding to their polar headgroups. Among 39 species, 22 species were detected to increase significantly in RSL3-treated Calu-1 by more than tenfold (Fig. 1c,d). The MS peak areas of 16 species increased progressively with RSL3 treatment time, and the increase of all 39 oxidized phospholipids was suppressed by Lip-1 (Fig. 1c).

**Table 1.**
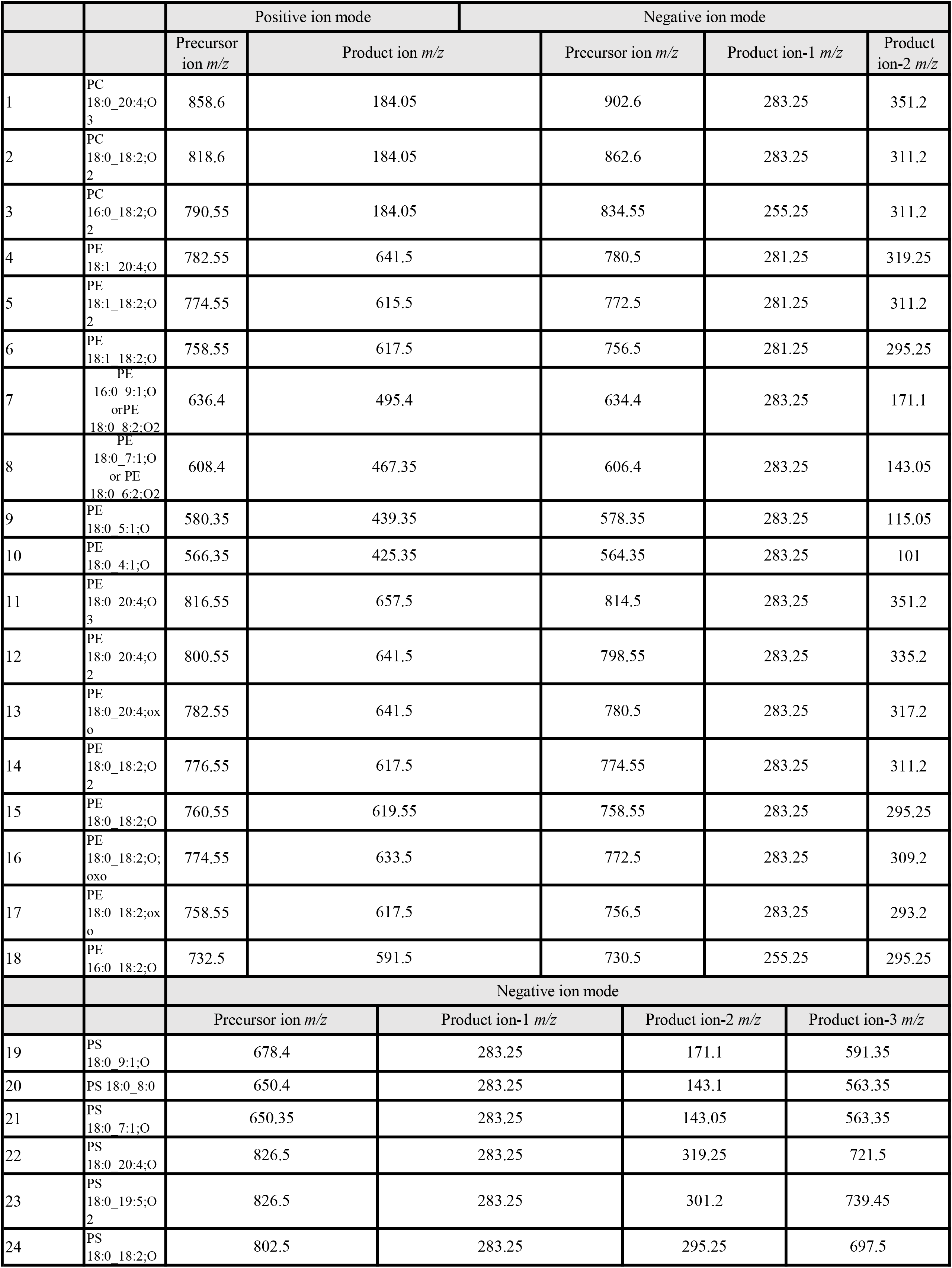
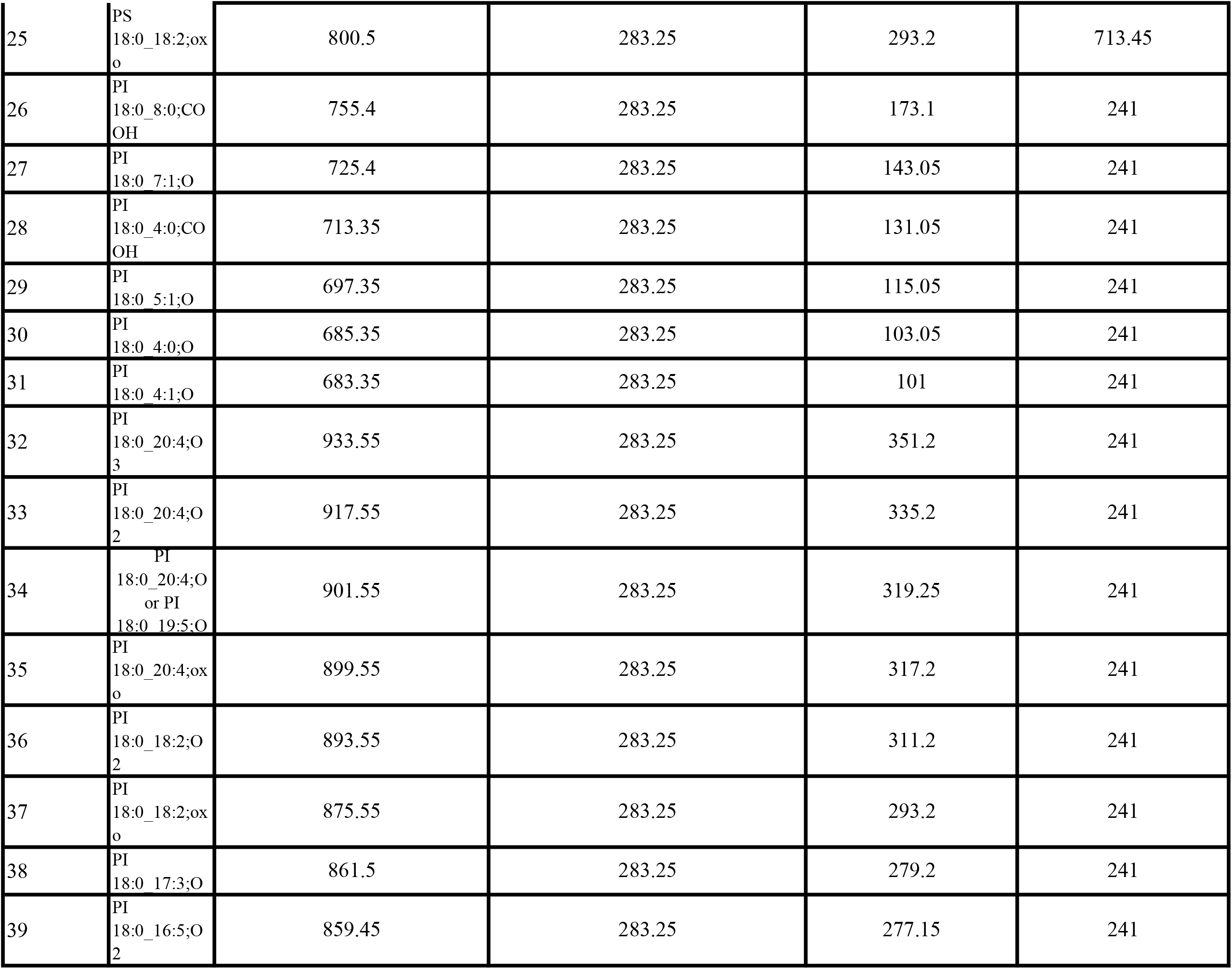
oxidized phospholipid species that increased in Calu-1 cells following RSL3 treatment.

|  |  | Positive ion mode |  | Negative ion mode |  |  |
| --- | --- | --- | --- | --- | --- | --- |
|  |  | Precursor ion *m/z* | Product ion *m/z* | Precursor ion *m/z* | Product ion-1 *m/z* | Product ion-2 *m/z* |
| 1 | PC<br>18:0_20:4;O<br>3 | 858.6 | 184.05 | 902.6 | 283.25 | 351.2 |
| 2 | PC<br>18:0_18:2;O<br>2 | 818.6 | 184.05 | 862.6 | 283.25 | 311.2 |
| 3 | PC<br>16:0_18:2;O<br>2 | 790.55 | 184.05 | 834.55 | 255.25 | 311.2 |
| 4 | PE<br>18:1_20:4;O | 782.55 | 641.5 | 780.5 | 281.25 | 319.25 |
| 5 | PE<br>18:1_18:2;O<br>2 | 774.55 | 615.5 | 772.5 | 281.25 | 311.2 |
| 6 | PE<br>18:1_18:2;O | 758.55 | 617.5 | 756.5 | 281.25 | 295.25 |
| 7 | PE<br>16:0_9:1;O<br>or PE<br>18:0_8:2;O2 | 636.4 | 495.4 | 634.4 | 283.25 | 171.1 |
| 8 | PE<br>18:0_7:1;O<br>or PE<br>18:0_6:2;O2 | 608.4 | 467.35 | 606.4 | 283.25 | 143.05 |
| 9 | PE<br>18:0_5:1;O | 580.35 | 439.35 | 578.35 | 283.25 | 115.05 |
| 10 | PE<br>18:0_4:1;O | 566.35 | 425.35 | 564.35 | 283.25 | 101 |
| 11 | PE<br>18:0_20:4;O<br>3 | 816.55 | 657.5 | 814.5 | 283.25 | 351.2 |
| 12 | PE<br>18:0_20:4;O<br>2 | 800.55 | 641.5 | 798.55 | 283.25 | 335.2 |
| 13 | PE<br>18:0_20:4;ox<br>o | 782.55 | 641.5 | 780.5 | 283.25 | 317.2 |
| 14 | PE<br>18:0_18:2;O<br>2 | 776.55 | 617.5 | 774.55 | 283.25 | 311.2 |
| 15 | PE<br>18:0_18:2;O | 760.55 | 619.55 | 758.55 | 283.25 | 295.25 |
| 16 | PE<br>18:0_18:2;O;<br>oxo | 774.55 | 633.5 | 772.5 | 283.25 | 309.2 |
| 17 | PE<br>18:0_18:2;ox<br>o | 758.55 | 617.5 | 756.5 | 283.25 | 293.2 |
| 18 | PE<br>16:0_18:2;O | 732.5 | 591.5 | 730.5 | 255.25 | 295.25 |
|  |  | Negative ion mode |  |  |  |  |
|  |  | Precursor ion *m/z* | Product ion-1 *m/z* | Product ion-2 *m/z* | Product ion-3 *m/z* |  |
| 19 | PS<br>18:0_9:1;O | 678.4 | 283.25 | 171.1 | 591.35 |  |
| 20 | PS 18:0_8:0 | 650.4 | 283.25 | 143.1 | 563.35 |  |
| 21 | PS<br>18:0_7:1;O | 650.35 | 283.25 | 143.05 | 563.35 |  |
| 22 | PS<br>18:0_20:4;O | 826.5 | 283.25 | 319.25 | 721.5 |  |
| 23 | PS<br>18:0_19:5;O<br>2 | 826.5 | 283.25 | 301.2 | 739.45 |  |
| 24 | PS<br>18:0_18:2;O | 802.5 | 283.25 | 295.25 | 697.5 |  |

|  |  |  |  |  |  |
| --- | --- | --- | --- | --- | --- |
| 25 | PS<br>18:0_18:2;ox<br>o | 800.5 | 283.25 | 293.2 | 713.45 |
| 26 | PI<br>18:0_8:0;CO<br>OH | 755.4 | 283.25 | 173.1 | 241 |
| 27 | PI<br>18:0_7:1;O | 725.4 | 283.25 | 143.05 | 241 |
| 28 | PI<br>18:0_4:0;CO<br>OH | 713.35 | 283.25 | 131.05 | 241 |
| 29 | PI<br>18:0_5:1;O | 697.35 | 283.25 | 115.05 | 241 |
| 30 | PI<br>18:0_4:0;O | 685.35 | 283.25 | 103.05 | 241 |
| 31 | PI<br>18:0_4:1;O | 683.35 | 283.25 | 101 | 241 |
| 32 | PI<br>18:0_20:4;O<br>3 | 933.55 | 283.25 | 351.2 | 241 |
| 33 | PI<br>18:0_20:4;O<br>2 | 917.55 | 283.25 | 335.2 | 241 |
| 34 | PI<br>18:0_20:4;O<br>or PI<br>18:0_19:5;O | 901.55 | 283.25 | 319.25 | 241 |
| 35 | PI<br>18:0_20:4;ox<br>o | 899.55 | 283.25 | 317.2 | 241 |
| 36 | PI<br>18:0_18:2;O<br>2 | 893.55 | 283.25 | 311.2 | 241 |
| 37 | PI<br>18:0_18:2;ox<br>o | 875.55 | 283.25 | 293.2 | 241 |
| 38 | PI<br>18:0_17:3;O | 861.5 | 283.25 | 279.2 | 241 |
| 39 | PI<br>18:0_16:5;O<br>2 | 859.45 | 283.25 | 277.15 | 241 |

We next investigated oxidized phospholipid profiles in Calu-1 cells treated with other ferroptosis inducers, including erastin, FINO2 and FIN56. Under each condition, multiple oxidized phospholipid species were also significantly increased by more than tenfold (Fig. 1e–g), similar to RSL3. In contrast, treatment of the ferroptosis-low-susceptible lung cancer cell lines H292 and A549^20^ with the same concentration and duration of RSL3 as in Calu-1 cells did not increase any oxidized phospholipid species by more than tenfold (Fig. 1h,i and Supplementary Fig. 1e).

Moreover, no oxidized phospholipid species increased by more than fivefold during apoptosis induced by staurosporine treatment (Supplementary Fig. 1f,g). Collectively, these results demonstrate that the increase of multiple oxidized phospholipid species is a characteristic feature of ferroptosis across ferroptosis-inducing stimuli but is not observed under ferroptosis-low-susceptible cell lines or other types of cell death like apoptosis.

### PE 18:0_20:4;O3 and PI 18:0_20:4;O3 are consistently increased across cellular and mouse models of ferroptosis

We next investigated whether specific oxidized phospholipids were consistently increased during ferroptosis across multiple ferroptosis-sensitive cell lines: human fibrosarcoma HT1080 cells, human lung cancer H661 cells, rat cardiomyoblast H9c2 cells, and the human pancreatic cancer cell lines MIAPaCa-2 and PANC-1. Consistent with the findings in Calu-1 cells, multiple oxidized phospholipid species increased more than tenfold in each cell line (Fig. 2a–e). Notably, PE 18:0_20:4;O3 and PI 18:0_20:4;O3 were the only two oxidized phospholipid species that increased significantly by more than tenfold in all six cell lines following RSL3 treatment (Fig. 2f) and in Calu-1 cells across four mechanistically distinct ferroptosis inducers (Fig. 1d–g).

**Fig. 2.**
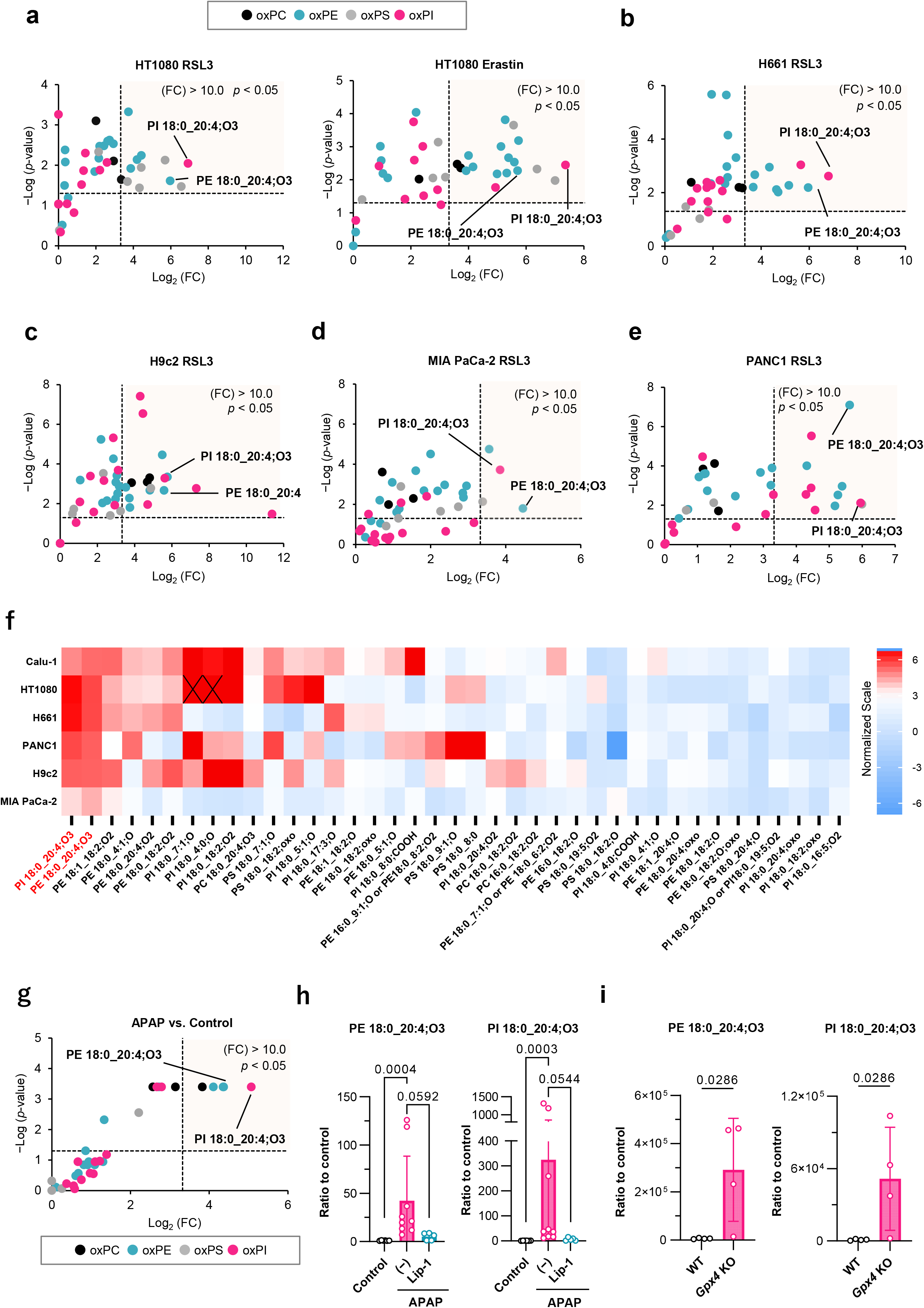
Oxidized phospholipid increase across diverse cell types and mouse models of ferroptosis. **a,** Fold changes in oxidized phospholipids in HT1080 cells treated with RSL3 (0.1 µM) or erastin (10 µM) for 5 h. **b–e,** Fold changes in oxidized phospholipids in H661 (**b**), H9c2 (**c**), MIA PaCa-2 (**d**) and PANC-1 (**e**) cells treated with RSL3 (0.1 µM) for 5 h. **f,** Number of oxidized phospholipid species that increased significantly by more than tenfold across six cell types, including Calu-1 cells. PE 18:0_20:4;O3 and PI 18:0_20:4;O3 increased in all six cell types examined. Crosses indicate no significant difference. Color scale indicates log2-transformed mean fold changes (**f**). **g,** Fold changes in oxidized phospholipids in liver tissues from APAP-treated mice (300 mg/kg, intraperitoneally, 4 h) compared with untreated mice. **h,** Fold changes in PE 18:0_20:4;O3 and PI 18:0_20:4;O3 in liver tissues from untreated mice, APAP-treated mice and mice treated with APAP followed by Lip-1 (10 mg/kg, intraperitoneally) 1 h after APAP administration. **i,** Analysis of oxidized phospholipids in the livers of *Alb-CreER^T^*^2^*;Gpx4^fl/fl^* mice. Data in (**a–f**) were obtained from three independent experiments, by the two-sided unpaired t-test. Data in (**g**) were obtained from n = 6–9 mice per group, by the Mann–Whitney U test. Data in (**h, i**) are mean ± s.d.; n = 5–9 mice per group (**h**) and 4 mice per group (**i**), by the Kruskal–Wallis test (**h**) or Mann–Whitney U test (**i**).

We next examined whether these oxidized phospholipids were also increased in two mouse models of ferroptosis-associated tissue injury: acetaminophen (APAP)-induced liver injury and a hepatocyte-specific *Gpx4* conditional knockout model. In the APAP-induced liver injury model^23^, hepatic injury, malondialdehyde accumulation, and histological damage were observed, all of which were suppressed by Lip-1 treatment (Supplementary Fig. 2a–c). Using this model, we quantified the 39 oxidized phospholipid species identified in Calu-1 cells. Seven oxidized phospholipid species, including PE 18:0_20:4;O3 and PI 18:0_20:4;O3, increased by more than tenfold, and these increases were markedly suppressed by Lip-1 treatment (Fig. 2g,h). To establish the hepatocyte-specific *Gpx4* conditional knockout model, we used *Gpx4*-floxed; *Alb-CreER^T^*^2^ mice that were fed a low-vitamin E diet following tamoxifen administration^24^. In this model, PE 18:0_20:4;O3 and PI 18:0_20:4;O3 were also significantly increased in liver tissues undergoing ferroptotic injury (Fig. 2i and Supplementary Fig. 2d). Collectively, these findings identify PE 18:0_20:4;O3 and PI 18:0_20:4;O3 as the two most consistently increased oxidized phospholipid species across six cultured cell lines and two mouse models of ferroptosis-associated tissue injury (Supplementary Fig. 1b).

### Structural characterization of PI 18:0_20:4;O3

We next sought to characterize the structure of PI 18:0_20:4;O3. LC–MS/MS analysis of lipid extracts from RSL3-treated Calu-1 cells revealed multiple fragment ions derived from PI 18:0_20:4;O3, including ions likely generated by cleavage of the oxidized arachidonoyl chain (FFA 20:4;O3) (Fig. 3a,b). These fragment ions were detected at several distinct retention times, indicating that PI 18:0_20:4;O3 comprises multiple structural isomers rather than a single molecular species.

**Fig. 3.**
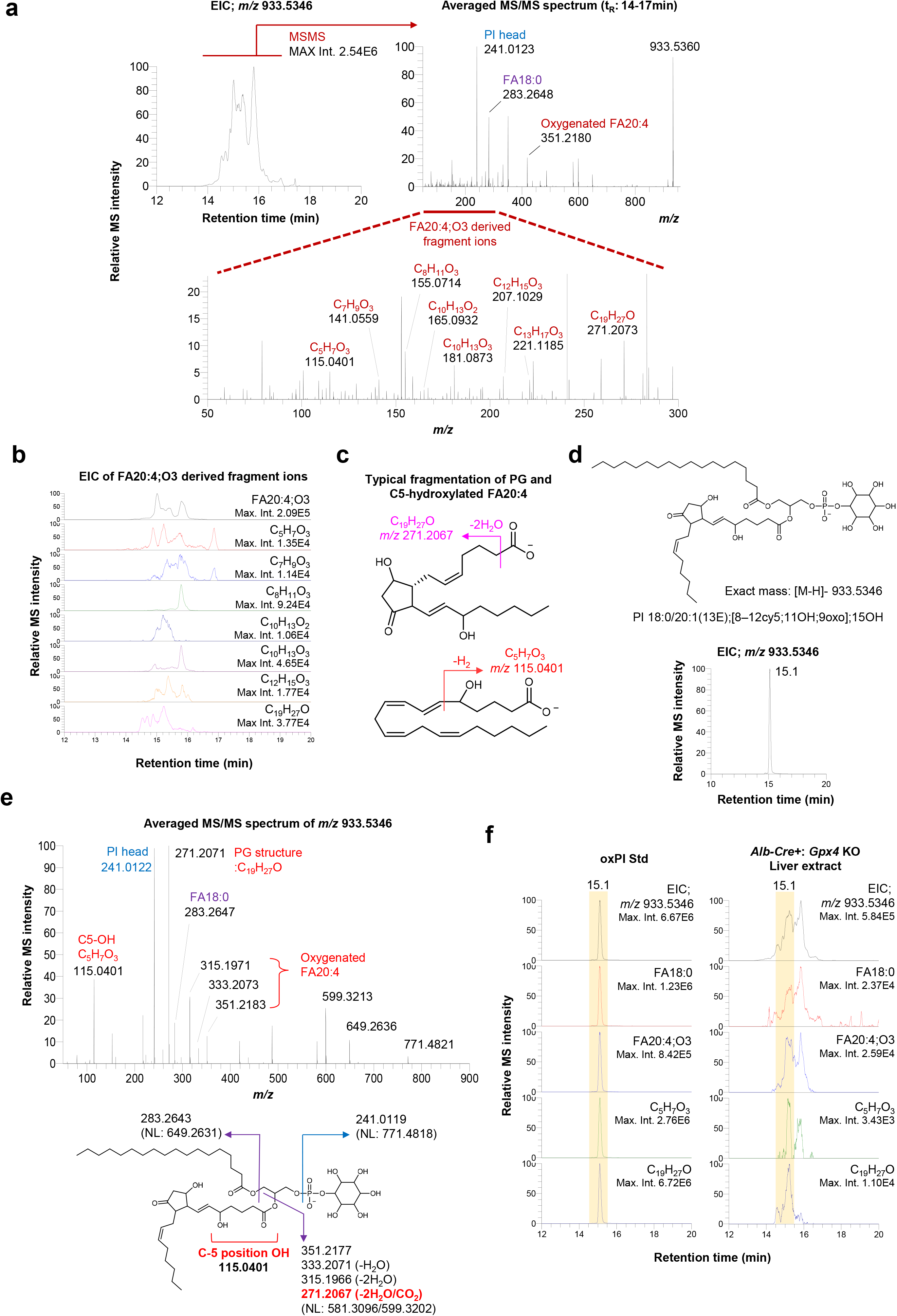
Structural characterization of oxidized phospholipid species commonly increased during ferroptosis. **a,** LC–MS/MS analysis of PI 18:0_20:4;O3 detected in lipid extracts from RSL3-treated Calu-1 cells. Fragment ions derived from the PI head group, FA18:0, and the oxygenated FA20:4 acyl chain are indicated. **b,** EICs of fragment ions derived from the oxidized arachidonoyl acyl chain (FA20:4;O3). **c,** Diagnostic fragment ions used for structural annotation of PI 18:0_20:4;O3. The C19H27O ion at *m/z* 271.2067 is characteristic of prostaglandin-like structures, whereas the C5H7O3 ion at *m/z* 115.0401 indicates oxidation at the C5 position of arachidonic acid. **d,e,** Chemical structure (**d**) and EICs and MS/MS spectra (**e**) of the synthetic PI 18:0/20:1(13E);[8– 12cy5;11OH;9oxo];15OH standard, an oxidized PI species containing a prostaglandin-like structure and a 5-hydroxy moiety. **f,** LC–MS/MS comparison of the synthetic PI 18:0/20:1(13E);[8– 12cy5;11OH;9oxo];15OH standard with the annotated PI 18:0_20:4;O3 species detected in liver extracts from *Alb-CreER^T^*^2^*;Gpx4^fl/fl^* mice.

We therefore focused on a relatively abundant fragment ion corresponding to C19H27O. In prostaglandins such as PGE2 and PGD2, a C19H27O-derived ion at *m/z* 271 has been reported as a major product ion ^25^. In addition, the prominent peak eluting at 15.1–15.2 min yielded a fragment ion corresponding to C5H7O3. This ion has been reported as a diagnostic fragment associated with oxidation at the C5 position of arachidonic acid derivatives, including 5-HETE ^26^ (Fig. 3c). Together, these fragmentation characteristics suggested that this chromatographic peak originated from an oxidized PI species containing an arachidonoyl chain with a prostaglandin-like cyclic structure and a hydroxyl group at the C5 position.

To further validate this structural assignment, we synthesized the corresponding authentic oxPI standard (Supplementary Fig. 3) and analyzed it by LC–MS/MS. The synthetic standard eluted at nearly the same retention time as the peak detected in RSL3-treated Calu-1 cells and exhibited a closely matching MS/MS fragmentation pattern (Fig. 3d,e). These results suggest that a subset of the PI 18:0_20:4;O3 species that increase in RSL3-treated Calu-1 cells corresponds to the structure shown in Fig. 3d. A species with the same chromatographic and MS/MS characteristics was also detected in liver tissue extracts from *Alb-CreER^T^*^2^*;Gpx4^fl/fl^*KO mice (Fig. 3f).

### Ferroptosis recruits PI4K2A to lysosomes but fails to activate PITT-mediated membrane repair

We next investigated whether the oxidized phospholipids consistently increase during the progression of ferroptosis. We previously demonstrated that LPO is initiated in lysosomes during the early stages of ferroptosis (Fig. 4a, left panel) ^20^. The resulting LMP allows labile iron to escape from lysosomes and promotes the propagation of LPO to the endoplasmic reticulum and plasma membrane, thereby accelerating ferroptosis. On the other hand, in the context of lysosomal damage, many reports indicate that LMP activates several lysosomal membrane-repair mechanisms^22, 27–29^.

**Fig. 4.**
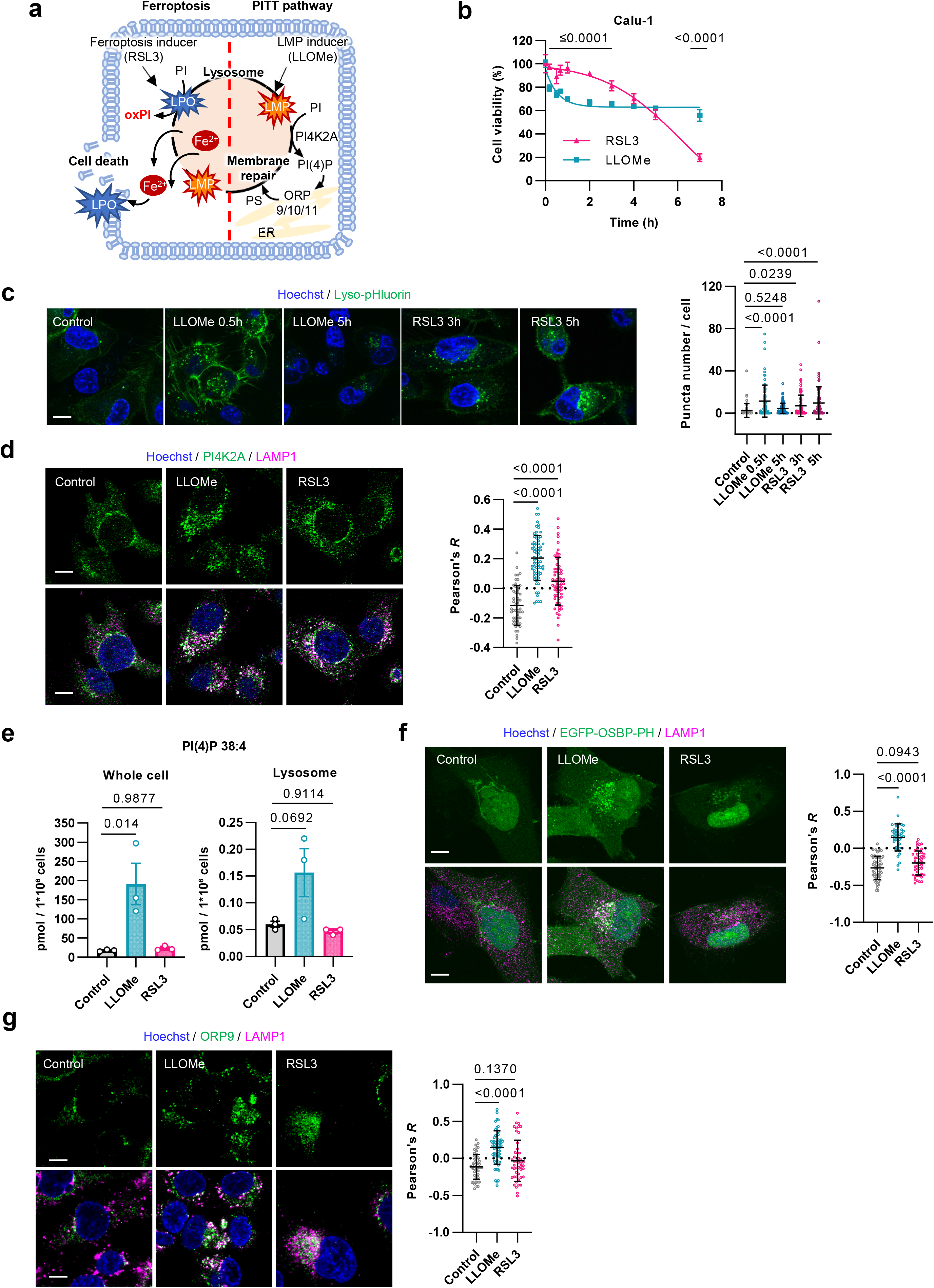
Ferroptosis impairs PITT-mediated lysosomal membrane repair. **a,** Schematic illustrations of the role of lysosomes as triggers of ferroptosis (left) and PITT-mediated lysosomal membrane repair following LMP (right). Created in BioRender. Yamada, K. (2027) https://BioRender.com/z5nmsby. **b,** Viability of Calu-1 cells treated with LLOMe (1 mM) or RSL3 (0.1 µM) for the indicated durations. **c,** Calu-1 cells stably expressing lyso-pHluorin were treated with LLOMe (1 mM) or RSL3 (0.1 µM) for the indicated durations. The number of lyso-pHluorin puncta per cell was quantified. **d,** Calu-1 cells were treated with LLOMe (1 mM) for 30 min or RSL3 (0.1 µM) for 3 h and immunostained for PI4K2A and LAMP1. **e,** LC–MS/MS analysis of PI(4)P 38:4 in whole-cell and lysosomal fractions. Calu-1 cells stably expressing TMEM192–3×HA were treated with LLOMe (1 mM) for 30 min or RSL3 (0.1 µM) for 3 h, followed by Lyso-IP. Values were normalized to cell number. **f,** Calu-1 cells stably expressing EGFP–OSBP–PH were treated with LLOMe (1 mM) or RSL3 (0.1 µM) for the indicated durations and immunostained for LAMP1. **g,** Calu-1 cells were treated with LLOMe (1 mM) for 30 min or RSL3 (0.1 µM) for 3 h and immunostained for ORP9 and LAMP1. Colocalization with LAMP1 was quantified using Pearson’s correlation coefficient (**d,f,g**). Data in (**b)** and (**e)** are presented as the mean ± s.d. of three independent experiments. Statistical significance was assessed using Sidak’s multiple-comparisons test (**b**) or the Tukey–Kramer test (**e**). Data in (**c,d)** and (**f,g)** are presented as the mean ± s.d. Each data point represents an individual cell quantified from three independent images; n = 88–111 (**c**), 52–69 (**d**), 42–53 (**f**), and 47–58 (**g**) cells. Statistical significance was assessed using Dunnett’s test. Scale bars, 10 µm (**c,d,f,g**).

The PITT pathway^22^ was recently identified as a lysosomal membrane-repair mechanism initiated by PI phosphorylation (Fig. 4a, right panel). In this pathway, Ca²⁺ released following LMP recruits phosphatidylinositol 4-kinase type 2 alpha (PI4K2A) to damaged lysosomes. PI4K2A subsequently phosphorylates PI at the 4-position to generate phosphatidylinositol 4-phosphate (PI(4)P) as a product. PI(4)P is then exchanged for endoplasmic reticulum-derived PS through the oxysterol-binding protein-related proteins ORP9, ORP10 and ORP11. This lipid exchange promotes lipid transport mediated by the autophagy-related protein ATG2, thereby facilitating lysosomal membrane repair.

Based on this evidence, we therefore hypothesized that although LMP occurs during ferroptosis, oxidation of PI, the substrate of PI4K2A, disrupts activation of the PITT pathway and thereby compromises lysosomal membrane repair (Fig. 4a). To test this hypothesis, we treated cells with RSL3 or the conventional LMP inducer L-leucyl-L-leucine methyl ester (LLOMe)^30^ and monitored cell viability over time. In Calu-1 and HT1080 cells, RSL3 treatment induced a time-dependent decrease in cell viability (Fig. 4b and Supplementary Fig. 4a). In contrast, LLOMe caused a rapid initial decrease in cell viability after the treatment, and then the cell viability remained at approximately 60–80% from 1 h onwards.

We next generated cells stably expressing the lysosome-targeted pH sensor Lyso-pHluorin^31^ and assessed lysosomal membrane damage and repair capacity by monitoring the lysosomal pH neutralization. LLOMe treatment markedly increased the number of Lyso-pHluorin-positive puncta after 30 min, indicating lysosomal membrane damage. However, puncta numbers returned to baseline by 5 h (Fig. 4c and Supplementary Fig. 4b). By contrast, Lyso-pHluorin-positive puncta progressively accumulated following RSL3 treatment and persisted for at least 5 h, indicating insufficient lysosomal membrane repair.

We then assessed the lysosomal recruitment of PI4K2A, an early event in the PITT pathway, by immunofluorescence microscopy. PI4K2A colocalized with the lysosomal membrane protein lysosome-associated membrane protein 1 (LAMP1) both 30 min after LLOMe treatment and 3 h after RSL3 treatment (Fig. 4d and Supplementary Fig. 4c). We subsequently quantified the contents of PI(4)P by LC–MS/MS. PI(4)P levels in Calu-1 cells increased markedly after 30 min of LLOMe treatment but remained largely unchanged after 3 h of RSL3 treatment compared to control cells (Fig. 4e). To examine the subcellular localization of PI(4)P, we generated Calu-1 and HT1080 cells stably expressing the fluorescent PI(4)P probe EGFP–OSBP-PH^32^. PI(4)P colocalized with lysosomes following LLOMe treatment, whereas no lysosomal colocalization was observed following RSL3 treatment (Fig. 4f). Similarly, ORP9, a downstream component of the PITT pathway, colocalized with lysosomes following LLOMe treatment but not following RSL3 treatment (Fig. 4g and Supplementary Fig. 4d).

Together, these results demonstrate that although PI4K2A is recruited to lysosomes during RSL3-induced ferroptosis in Calu-1 and HT1080 cells, the downstream PITT pathway is not efficiently activated.

### Oxidized PIs inhibit PI4K2A-dependent PI(4)P production and disrupt lysosomal repair during ferroptosis

We next investigated the mechanistic relationship between increased oxPIs and the failure of PITT pathway activation during ferroptosis. Oxidized phospholipid profiling was analyzed by LC-MS/MS in lysosomes isolated from Calu-1 cells treated with RSL3 or LLOMe. PI 18:0_20:4;O3 was significantly increased in the isolated lysosomal fraction following RSL3 but not LLOMe treatment (Fig. 5a and Supplementary Fig. 5a).

**Fig. 5.**
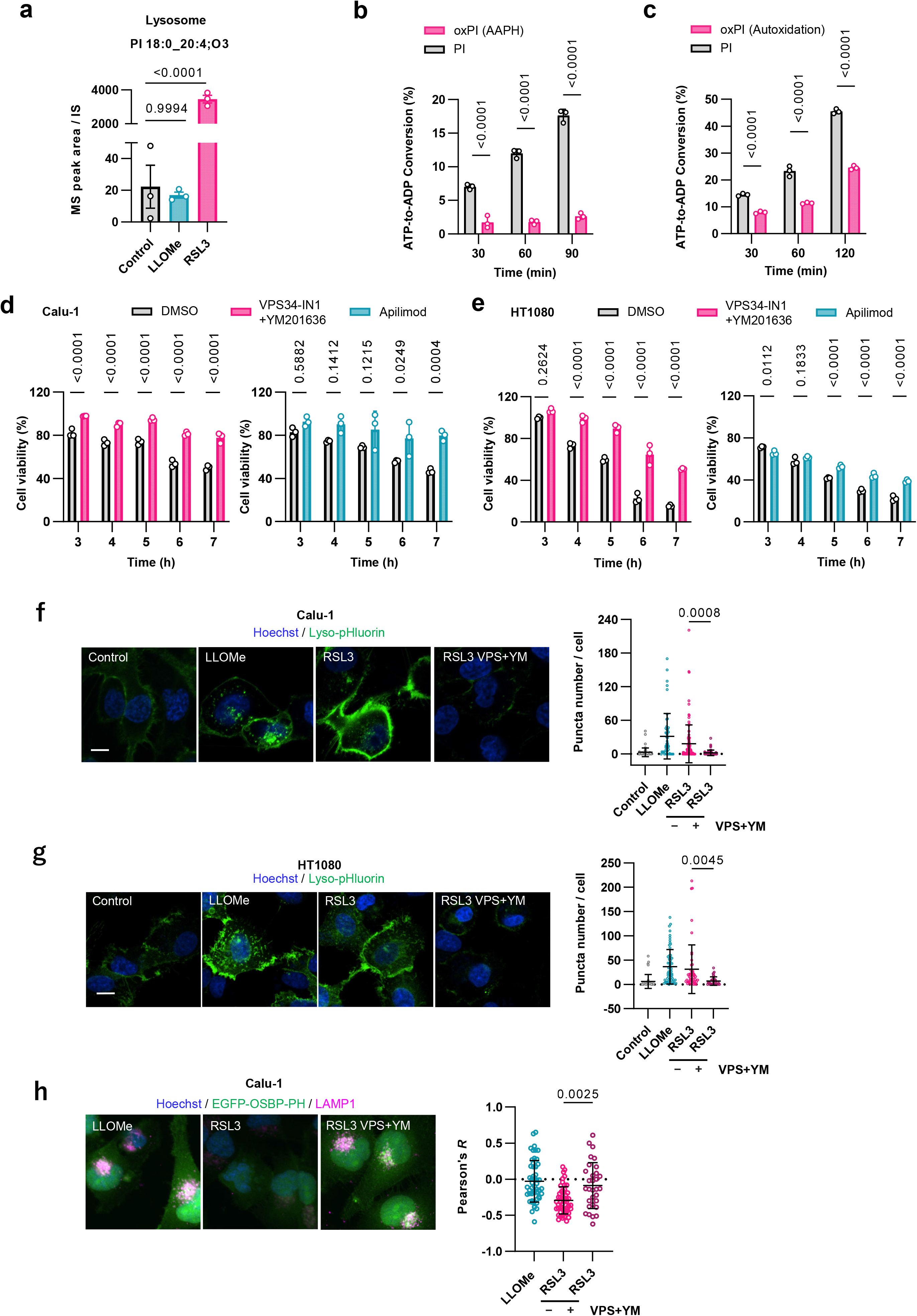
oxPIs inhibit PI4K2A-dependent PI(4)P production and disrupt lysosomal membrane repair during ferroptosis. **a,** LC–MS/MS-based lysosomal lipidomic analysis. Calu-1 cells stably expressing TMEM192– 3×HA were treated with LLOMe (1 mM, 30 min) or RSL3 (0.1 µM, 3 h), followed by Lyso-IP. PI 18:0_20:4;O3 in lysosomal fractions were quantified and normalized to the internal standard. **b,c,** PI4K2A activity assays using recombinant PI4K2A (10 ng per well) and SAPI (0.2 µg per well) for the indicated durations. Non-oxidized SAPI was compared with AAPH-oxidized SAPI in (**b)** and autoxidized SAPI in (**c)**. Kinase activity was measured using the ADP-Glo Kinase Assay. **d,e,** Viability of Calu-1 (**d**) and HT1080 (**e**) cells pretreated with a combination of VPS34-IN1 and YM201636 (1 µM each) or apilimod (1 µM), followed by treatment with RSL3 (0.1 µM) for the indicated durations. **f,g,** Calu-1 (f) and HT1080 (g) cells stably expressing lyso-pHluorin were pretreated for 24 h with or without a combination of VPS34-IN1 and YM201636 (1 µM each), followed by treatment with LLOMe (1 mM, 30 min) or RSL3 (0.1 µM, 3 h). The number of lyso-pHluorin puncta per cell was quantified. **h,** Calu-1 cells stably expressing EGFP–OSBP–PH were pretreated for 24 h with or without a combination of VPS34-IN1 and YM201636 (1 µM each), followed by treatment with LLOMe (1 mM, 30 min) or RSL3 (0.1 µM, 3 h). Colocalization of EGFP–OSBP–PH with LAMP1 was quantified using Pearson’s correlation coefficient in (**h)**. Data in (**a–e)** are presented as the mean ± s.d. of three independent experiments. Statistical significance was assessed using Dunnett’s test (**a**) or Sidak’s multiple-comparisons test (**b–e**). Data in (**f–h)** are presented as the mean ± s.d. Each data point represents an individual cell quantified from three independent images; n = 51-101 (**f**), 37–62 (**g**), and 31-52 (**h**) cells. Statistical significance was assessed using the Tukey–Kramer test. Scale bars, 10 µm (**f–h**).

We then used an ADP-Glo kinase assay to determine whether oxPI could serve as a substrate for PI4K2A. PI4K2A exhibited robust kinase activity when non-oxidized SAPI (or PI 18:0_20:4) was used as the substrate. In contrast, only minimal PI4K2A-dependent kinase activity was detected when AAPH-oxidized SAPI was used (Fig. 5b). To more closely approximate the intracellular lipid environment, PI4K2A activity was also measured in the presence of both oxidized- and non-oxidized PI (Supplementary Fig. 5b). Even under these conditions, the presence of oxPIs significantly reduced PI4K2A-dependent conversion of ATP to ADP (Fig. 5c), suggesting that oxPIs may interfere PI4K2A-dependent kinase activity.

Post-translational modifications also regulate PI4K2A activity. Palmitoylation promotes stable membrane association and enhances kinase activity^33^, whereas the phosphorylation of PI4K2A regulates protein interactions and intracellular trafficking^34^. However, no significant differences in PI4K2A palmitoylation (Supplementary Fig. 5c) or phosphorylation (Supplementary Fig. 5d) were observed between LLOMe- and RSL3-treated cells. These results support a model in which oxPIs generated during ferroptosis impair PI(4)P production, thereby compromising PITT-mediated lysosomal membrane repair.

We next examined whether defective PITT pathway activation contributes to ferroptotic cell death. We hypothesized that increasing PI(4)P levels would suppress ferroptosis. Lysosomal PI(4)P has been reported to accumulate following inhibition of the PI(3)P–PI(3,5)P₂ synthesis pathway mediated by phosphatidylinositol 3-kinase catalytic subunit type 3 (PIK3C3) and phosphoinositide kinase, FYVE-type zinc finger containing (PIKfyve)^35^. We therefore assessed cell viability during ferroptosis following combined treatment with the PIK3C3 inhibitor VPS34-IN1 and the PIKfyve inhibitor YM201636, or following pretreatment with the selective PIKfyve inhibitor apilimod (Supplementary Fig. 5e). Both combined VPS34-IN1 and YM201636 treatment and apilimod pretreatment attenuated RSL3-induced cell death (Fig. 5d,e). Consistent with these findings, analysis of cells stably expressing Lyso-pHluorin showed that both combined VPS34-IN1 and YM201636 treatment and apilimod pretreatment suppressed LMP in Calu-1 and HT1080 cells (Fig. 5f,g). Also, VPS34-IN1 and YM201636 combined treatment increased PI(4)P colocalized with lysosomes (Fig. 5h). Collectively, these results suggest that maintaining sufficient PI(4)P levels at lysosomes enables activation of the PITT pathway and thereby suppresses ferroptosis.

## Discussion

In this study, we identified PE 18:0_20:4;O3 and PI 18:0_20:4;O3 as two recurrent oxidized phospholipid species that consistently increased during ferroptosis across multiple cell types and induction conditions. Previous studies have extensively documented the increase of oxidized phospholipids, particularly oxPEs, during ferroptosis. Kagan and colleagues identified four oxPEs, including PE 18:0_20:4;O3, in Pfa1 cells undergoing ferroptosis following RSL3 treatment or GPX4 deficiency, and established 15-HpETE-PE as a representative ferroptosis-associated lipid^10, 36^. Other groups have similarly reported increased PE 18:0_20:4;O3 abundance during ferroptosis in RSL3-treated MDA-MB-231 and Huh-7 cells^37, 38^. Consistent with these findings, we observed increased PE 18:0_20:4;O3 in six distinct cultured cell lines following treatment with multiple ferroptosis inducers, as well as in two mouse models. Thus, our findings extend previous observations by showing that PE 18:0_20:4;O3 is reproducibly increased across diverse experimental models.

Although oxPEs have received considerable attention in ferroptosis research, the biological significance of oxPIs remains poorly understood. Kagan and colleagues reported that PI 18:0_20:4;O3 increased following ferroptosis induction in Pfa1 cells^10^. However, because the abundance of its non-oxidized precursor, PI 18:0_20:4, was not reduced in acyl-CoA synthetase long chain family member 4 (ACSL4)-deficient cells, PI 18:0_20:4;O3 was not considered a characteristic ferroptosis-associated oxidized phospholipid. By contrast, Doll and colleagues reported reduced PI 18:0_20:4 abundance in ACSL4-deficient Pfa1 cells^11^. Increased PI 18:0_20:4;O3 has also been observed during ferroptosis in bone marrow-derived macrophages and MDA-MB-231 cells^13, 37^. Together with our findings, these observations suggest that PI 18:0_20:4;O3 is not restricted to a particular cell type or induction condition, but represents a reproducibly increased ferroptosis-associated oxidized phospholipid across distinct experimental systems.

Structural analysis revealed that PI 18:0_20:4;O3 comprises multiple structural isomers rather than a single oxidation product. This is consistent with the complex oxidation chemistry of arachidonic acid, in which free radical-mediated oxidation can generate a diverse array of positional and structural isomers. Among these isomers, we identified a PI species containing an oxidized arachidonoyl chain characterized by a prostaglandin-like cyclic structure and oxidation at the C5 position. Comparison with a synthetic standard supported this structural assignment, and the same species was detected in both RSL3-treated Calu-1 cells and the livers of *Alb-CreER^T^*^2^*;Gpx4^fl/fl^* KO mice. These findings indicate that structurally complex oxPI species are generated in association with ferroptosis both *in vitro* and *in vivo*.

Previous studies of ferroptosis-associated phospholipid oxidation have predominantly focused on hydroperoxy-PE species containing arachidonic or adrenic acid^10^. In contrast, the detailed molecular structures of oxPI species generated during ferroptosis remain largely unexplored. Our findings therefore expand the known structural diversity of ferroptosis-associated oxidized phospholipids and reveal that oxPI is generated as a heterogeneous population of structurally distinct oxidation products rather than as a single uniform species.

Free radical-mediated oxidation of arachidonic acid is known to generate isoprostanes containing prostaglandin-like cyclic structures^39–41^. Importantly, such oxidation can occur while arachidonic acid remains esterified within membrane phospholipids, resulting in the formation of structurally complex oxidized phospholipids, including esterified isoprostanes^40, 41^. Our findings suggest that similar free radical-mediated chemistry occurs during ferroptosis-associated membrane LPO, leading to the generation of structurally complex oxPI species.

Although the retention time and MS/MS fragmentation pattern of the endogenous lipid closely matched those of the synthetic standard, these analyses alone cannot definitively determine all double-bond positions or the stereochemical configuration of the endogenous species. More comprehensive structural analyses, including approaches capable of resolving double-bond positions, such as oxygen attachment dissociation (OAD)-MS/MS, together with isolation and detailed characterization of the target oxidized lipid from biological samples, will therefore be required to establish its complete molecular structure. Such analyses represent an important direction for future studies.

Given the consistent increase of oxPIs during ferroptosis, we focused on the PITT pathway, a lysosomal membrane repair system that depends on PI as a substrate. Under our experimental conditions, PI(4)P-dependent lysosomal membrane repair was impaired during ferroptosis, whereas PI(4)P supplementation partially suppressed ferroptotic cell death. Although Zhang and colleagues did not directly examine the PITT pathway or PI(4)P, they reported that overexpression of OSBPL10 (ORP10) promoted PS-dependent lysosomal membrane repair and attenuated LMP and ferroptosis in neurons following spinal cord injury^42^. Similarly, Li and colleagues demonstrated that lysophosphatidylcholine acyltransferase 1 (LPCAT1) overexpression increased endoplasmic reticulum–lysosome contacts and phospholipid delivery in nucleus pulposus cells, thereby suppressing LMP and ferroptosis through enhanced lysosomal membrane repair^43^. More recently, a tectonin beta-propeller repeat containing 1 (TECPR1)-mediated pathway was identified as a PI(4)P-dependent lysosomal membrane repair mechanism distinct from the PITT pathway^44^. In our study, PI(4)P failed to accumulate on lysosomal membranes during ferroptosis, raising the possibility that TECPR1-mediated repair was also insufficiently activated under these conditions. Nevertheless, other PI(4)P-independent lysosomal membrane repair mechanisms, including ESCRT- and sphingomyelin-mediated pathways, also exist^27, 28, 45^, and their involvement in ferroptosis has been proposed^46^. Collectively, our findings suggest that reduced PI(4)P production during ferroptosis impairs PITT-mediated lysosomal membrane repair, further supporting a central role for lysosomes in determining ferroptosis susceptibility.

We further found that PI4K2A accumulated on lysosomes during ferroptosis, despite a significant reduction in PI(4)P production. Moreover, oxPIs did not appear to serve as a substrate for PI4K2A. However, because most PI molecules in biological membranes likely remain unoxidized, replacing only a small fraction of the PI pool with oxPIs may not fully account for the failure of lysosomal membrane repair. We therefore considered several mechanisms that could link oxPIs formation to disruption of lysosomal membrane repair.

LPO chain reactions are likely to occur in the immediate vicinity of ferroptosis-associated lysosomal membrane lesions. Thus, even in the minimal amount of oxidized phospholipids in the membrane fraction, their local abundance at sites of lysosomal membrane damage may be substantially increased. Oxidized phospholipids can markedly alter membrane biophysical properties by reducing membrane thickness, increasing the area per lipid molecule, and promoting membrane curvature^47, 48^. Their local generation at sites of lysosomal membrane damage may therefore alter membrane organization and change the orientation, distribution, or accessibility of PI at the membrane surface. In addition, oxidation of PI itself would be expected to alter its intramembrane conformation and topology, as proposed by the lipid whisker model^49, 50^.

PI4K2A is a membrane-associated PI 4-kinase for which an ADP-bound X-ray crystal structure has been reported^51^. However, the structure of PI4K2A bound to PI has not yet been resolved, and its substrate-binding mode remains unclear. PI4K2A activity is regulated by membrane cholesterol, which modulates membrane fluidity and fluctuations, suggesting that an appropriate membrane environment is required to sustain efficient catalytic activity^51^. PI is also predicted to bind PI4K2A by inserting its glycerol backbone and inositol headgroup into the substrate-binding pocket while retaining its fatty acyl chains within the membrane.

Taken together, these considerations suggest that oxidized phospholipids generated at sites of lysosomal membrane damage during ferroptosis perturb the membrane environment required for efficient PI4K2A activity. Such changes may affect the membrane-associated configuration of PI4K2A and productive engagement of PI with the PI4K2A substrate-binding pocket. Consequently, despite its recruitment to lysosomes, PI4K2A may be unable to efficiently recognize and phosphorylate PI, resulting in insufficient PI(4)P production and impaired PITT-mediated membrane repair.

In addition to membrane repair, the recruitment of antioxidant enzymes to lysosomes may reinforce local defense mechanisms and thereby influence ferroptosis susceptibility. For example, in lymph node-metastatic melanoma cells, lysosomal accumulation of FSP1 has been reported to suppress LPO and confer resistance to ferroptosis^52^. This observation further underscores the importance of lysosomes in regulating ferroptotic cell death.

In conclusion, our findings suggest that oxPIs generated during ferroptosis contribute to dysfunction of the PITT-mediated lysosomal membrane repair pathway. Considering ferroptosis as a collapse of organelle-centered cellular resilience may provide a useful framework for understanding how individual organelles sense, withstand, and repair oxidative lipid damage. Defining these organelle-specific protective mechanisms may deepen our understanding of ferroptosis and ultimately facilitate the development of therapeutic strategies for neurodegenerative diseases and cancer.

## Methods

### Chemicals and reagents

Unless otherwise specified, reagents and solvents were purchased from FUJIFILM Wako Pure Chemical Corporation or Nacalai Tesque. The following reagents were used in this study: RSL3 (Sigma-Aldrich, SML2234), liproxstatin-1 (Lip-1; Sigma-Aldrich, SML1414), acetaminophen (APAP; Sigma-Aldrich, A7085), erastin (Cayman Chemical, 17754), FINO2 (Cayman Chemical, 25096), FIN56 (Cayman Chemical, 25180), L-leucyl-L-leucine methyl ester (LLOMe; Cayman Chemical, 16008), 2-bromopalmitate (2-BP; BLDpharm, BD-16513), apilimod (Selleck Chemicals, S6414), VPS34-IN1 (Selleck Chemicals, S7980), YM201636 (Selleck Chemicals, S1219), staurosporine (Cayman Chemical, 81590), 1-stearoyl-2-arachidonoyl-sn-glycero-3-phosphoinositol (SAPI; Avanti Polar Lipids, 850144P), 2,2′-azobis(2-methylpropionamidine) dihydrochloride (AAPH; FUJIFILM Wako Pure Chemical Corporation, 017-21332), and fetal bovine serum (FBS; NICHIREI BIOSCIENCES, 175012).

### Cell culture

Calu-1 (HTB-54), H661 (HTB-183) and H292 (CRL-1848) were purchased from the ATCC; A549 (86012804) was from ECACC; HT1080 (IFO50354) was from the Japanese Collection of Research Bioresources (JCRB); PANC-1 (RCB2095) and MIA PaCa-2 (RCB2094) were from the RIKEN BioResource Research Center (BRC); Lenti-X 293T (632180) was from Takara Bio; and H9c2 cells were provided by the Department of Cardiovascular Medicine, Graduate School of Medical Sciences, Kyushu University.Calu-1 cells were cultured in McCoy’s 5A medium (Gibco). H661 cells were cultured in RPMI-1640 medium (FUJIFILM Wako Pure Chemical Corporation), H292 and PANC-1 cells were cultured in RPMI-1640 medium (Nacalai Tesque), H9c2, A549 and MIA PaCa-2 cells were cultured in low-glucose DMEM (Nacalai Tesque), Lenti-X 293T cells were cultured in high-glucose DMEM containing 4.5 g/L glucose (Nacalai Tesque), and HT1080 cells were cultured in MEM (Nacalai Tesque) supplemented with 1 mM sodium pyruvate (Sigma-Aldrich) and 0.1 mM MEM non-essential amino acids (Nacalai Tesque). All culture media were supplemented with 10% FBS, 100 U/mL penicillin, and 100 µg/mL streptomycin (Nacalai Tesque). All cell lines were grown at 37 °C in a humid atmosphere with 5% CO_2_.

### Cell viability assay

Cell viability was assessed using the MTT assay. Cells were seeded in 96-well plates at a density of 1 × 10^4^ cells per well and cultured for 24 h before treatment with the indicated compounds. For time-course experiments, cells were treated for the indicated durations, whereas concentration-dependent experiments were performed using a 24-h treatment period. MTT solution (Nacalai Tesque) was added to each well to a final concentration of 5 mg/mL, and the cells were incubated for 1 h at 37 °C in 5% CO2. The medium was removed, and 100 µl DMSO was added to each well. Absorbance at 570 and 650 nm was measured using an EnSpire Multimode Plate Reader (PerkinElmer) or an ARVO Kira multimode plate reader (Revvity). Background-corrected absorbance was calculated by subtracting the absorbance at 650 nm from that at 570 nm. Relative cell viability was calculated by setting the absorbance of the control group to 100%.

### LDH release assay

Cells were seeded in 96-well plates at a density of 1 × 10^4^ cells per well and cultured for 24 h before treatment. LDH release was measured using the Cytotoxicity LDH Assay Kit-WST (Dojindo) according to the manufacturer’s instructions. After treatment, 100 µl culture supernatant was transferred to a new 96-well plate and mixed with the assay buffer and detection reagent. After incubation for 30 min at room temperature, stop solution was added, and absorbance at 490 nm was measured using an EnSpire Multimode Plate Reader (PerkinElmer). LDH release was calculated by setting the absorbance of cells treated with lysis buffer to 100%.

### Flow cytometric analysis

Cells were seeded in six-well plates at a density of 2 × 10^5^ cells per well and cultured for 24 h. After treatment with RSL3 for 5 h, cells were incubated with C11-BODIPY for 20 min at 37 °C in 5% CO2. The medium was removed, and the cells were washed with 1 mL PBS and detached using 1 mL Accutase for 5 min. Cells were collected in 2-mL tubes and centrifuged at 800 × g for 5 min. The pellets were washed once with 1 mL PBS, centrifuged again at 800 × g for 5 min, and resuspended in 0.5 mL HBSS. C11-BODIPY fluorescence was measured immediately using the FITC detector (511–543 nm). At least 10,000 events were acquired per sample, and the data were analyzed using FlowJo 10 software (Becton Dickinson).

### Lipid extraction from cultured cells

Cells were seeded in 10-cm dishes at a density of 1 × 10^6^ cells per dish and cultured for 24 h before treatment. Cells were treated with RSL3 (0.1 µM, 0.01% DMSO) for 5 h, erastin (10 µM, 0.01% DMSO) for 7 h, FINO2 (10 µM, 0.01% DMSO) for 6 h, or FIN56 (5 µM, 0.01% DMSO) for 5 h at 37 °C in 5% CO2. After treatment, 2 mL PBS was added, and the cells were harvested using a cell scraper. The cell suspensions were centrifuged at 15,000 rpm for 10 min, and the supernatants were removed. Each pellet was resuspended in 250 µl methanol precooled to −30 °C and containing 100 µM butylated hydroxytoluene (BHT) and 100 µM EDTA. The suspension was sonicated using a Sonifier 250D (Branson) at an output setting of 1.5 for ten 1-s pulses separated by 1-s intervals.

Samples were centrifuged at 15,000 rpm for 10 min, and the resulting supernatants were collected as lipid extracts. Samples were stored at −80 °C and analyzed by LC–MS/MS within 24 h.

### Apoptosis induction and analysis

Apoptosis was induced in HT1080 cells by treatment with staurosporine (STS; 0.1 µM) for 24 h, followed by treatment with STS (1 µM) for an additional 4 h. Apoptotic cells were detected using the Annexin V-FITC Apoptosis Detection Kit (Nacalai Tesque, 15342-54) according to the manufacturer’s instructions. Fluorescence was measured using a FACSVerse flow cytometer (Becton Dickinson). Annexin V–FITC was excited at 488 nm, and emission was detected using a 527/32-nm filter. Propidium iodide was excited at 488 nm, and emission was detected using a 700/54-nm filter.

### Lentiviral production

Lenti-X 293T cells (3 × 10^5^ cells/well) were seeded in 12-well plates and cultured overnight. Subsequently, 0.4 µg of envelope plasmid pMD2.G (Addgene, 12259), 0.6 µg of packaging plasmid psPAX2 (Addgene, 12260), 0.8 µg of cloned lentiviral plasmid, and 10.8 µg of PEI MAX (Polysciences, 24765-100) were mixed in 200 µL of Opti-MEM (Gibco, 31985062). The solution was added dropwise to Lenti-X 293T cells cultured in 300 µL of medium. After 5 h, the medium was replaced, and the cells were cultured for additional 2 days. Culture supernatants were collected and centrifuged at 2,000 × g for 10 min, and the resulting supernatants were used as viral stocks.

Plasmids encoding EGFP–OSBP–PH and lyso-pHluorin were generated by inserting the corresponding coding sequences into linearized pLJM1-FLAG-GFP-TMEM192 (Addgene, 134630). pLJM1-TMEM192-mRFP-3×HA (Addgene, 134631) was used to generate cells for Lyso-IP. All plasmids used in this study are listed in Table 2.

**Table 2.** Plasmids used in this study.

| Plasmid | Source |
| --- | --- |
| EGFP-OSBP-PH | Addgene no. 49571 |
| pCMV-lyso-pHluorin | Addgene no. 70113 |
| pLJM1-FLAG-GFP-TMEM192 | Addgene no. 134630 |
| pLJM1-TMEM192-mRFP-3×HA | Addgene no. 134631 |
| pMD2.G | Addgene no. 12259 |
| psPAX2 | Addgene no. 12260 |

### Generation of stable cell lines

Calu-1 or HT1080 cells were seeded in 12-well plates at a density of 2.5 × 10^4^ cells per well and cultured overnight. The medium was replaced with 400 µL of fresh medium containing 10 µg/mL polybrene (Nacalai Tesque, 129661-81), and 100 µL of viral supernatant was added. After 24 h, the medium was replaced with fresh medium, and the cells were cultured for an additional 48 h. Stable cell populations were selected using puromycin (5 µg/mL; Nacalai Tesque, 19752-64) for 3 days.

### Live-cell confocal imaging and image analysis

Cells were seeded in four-well CELLview glass-bottom dishes (Greiner Bio-One) at a density of 3 × 10^4^ cells per compartment and cultured for 24 h. After treatment and staining, cells were washed three times with PBS. Images were acquired using either an LSM 700 confocal laser-scanning microscope (Carl Zeiss Microscopy GmbH) with a ×40 objective lens and ZEN 2011 v14 software (Carl Zeiss), or an AX confocal microscope (Nikon) with a ×40 objective lens and NIS-Elements AR 6.20.00 software (Nikon). Images were processed using ZEN 3.5 software (Carl Zeiss Microscopy GmbH) or NIS-Elements Viewer 5.22.00 software (Nikon), respectively.

The number of puncta and mean fluorescence intensity per cell were quantified using Fiji (ImageJ) 1.54 software (NIH). Cell boundaries were manually defined as regions of interest, and three independent images were analyzed for each group. The total number of cells analyzed across the three images was defined as n. Colocalization was quantified using the Coloc 2 in ImageJ with Pearson’s correlation coefficient (Pearson’s r).

### Immunofluorescence imaging

Cells were seeded in four-well CELLview glass-bottom dishes (Greiner Bio-One) at a density of 3 × 10^4^ cells per compartment and cultured for 24 h. After treatment, cells were washed three times with PBS and fixed with 4% paraformaldehyde for 10 min at room temperature. Cells were permeabilized and blocked for 30 min at room temperature using 5% normal donkey serum (Jackson ImmunoResearch, 017-000-121) in PBST containing 0.3% Triton X-100.

Cells were incubated overnight at 4 °C with mouse anti-PI4K2A antibody (Santa Cruz Biotechnology, sc-390026; 1:200), mouse anti-ORP9 antibody (Santa Cruz Biotechnology, sc-398961; 1:200) or rabbit anti-LAMP1 antibody (Cell Signaling Technology, 9091; 1:200), as indicated. After three washes with PBST, cells were incubated for 1 h at room temperature with Alexa Fluor 488-conjugated donkey anti-mouse IgG (Jackson ImmunoResearch, 715-545-151; 1:500), Alexa Fluor 647-conjugated donkey anti-rabbit IgG (Jackson ImmunoResearch, 711-605-152; 1:500) and Hoechst 33342 (1 µg/mL; Nacalai Tesque, 19172-51). After three washes with PBST, images were acquired using either an LSM 700 confocal laser-scanning microscope (Carl Zeiss Microscopy GmbH) with a ×40 objective lens, or an AX confocal microscope (Nikon) with a ×40 objective lens.

Excitation and emission settings were as follows: Hoechst33342, λex =405 nm, λem = 410–476 nm; Alexa Fluor 488, λex = 488 nm, λem = 516–559 nm; and Alexa Fluor 647, λex = 633 nm, λem = 638– 746 nm. Colocalization with LAMP1 was quantified using the Coloc 2 in ImageJ with Pearson’s correlation coefficient (Pearson’s r). Three independent images were analyzed for each group, and the total number of cells analyzed was defined as n.

### Lysosome immunoprecipitation

Lysosomes were isolated by lysosome immunoprecipitation (Lyso-IP^53, 54^) using anti-HA magnetic beads. Cells stably expressing TMEM192–3×HA were seeded in 15-cm dishes at a density of 4 × 10^6^ cells per dish and cultured for 24 h. After treatment, the cells were washed with PBS, harvested by scraping, and pelleted by centrifugation. Cell pellets were resuspended in 2 mL KPBS buffer and gently homogenized with 20 strokes using a Dounce homogenizer. Homogenates were centrifuged at 1,000 × g for 2 min, and the resulting supernatants were incubated with 100 µL anti-HA magnetic beads (Thermo Scientific, 88837) for 20 min. Lysosome-bound beads were washed three times with KPBS buffer using a DynaMag Spin Magnet (Thermo Scientific, 12320D).

For protein analysis, the beads were incubated with 100 µL lysis buffer for 10 min, and the eluates were collected as lysosomal fractions. Protein concentrations were determined using a Protein Assay BCA Kit (Nacalai Tesque). For lipid analysis, lipids were extracted from the beads using 200 µL ice-cold methanol containing 100 µM BHT and100 nM LysoPC 18:1(d7) (Avanti Polar Lipids, 791643) as an internal standard.

### Preparation of oxidized SAPI

For autoxidation, SAPI (1 mM, 100 µL) was transferred to a 1.5-mL tube and dried under a stream of nitrogen. The dried SAPI was incubated under an oxygen atmosphere at 37 °C for 6 d and subsequently redissolved in methanol.

For AAPH-induced oxidation, SAPI and AAPH were mixed in PBS at final concentrations of 55.3 µM and 5 mM, respectively, in a total volume of 1 mL. After vortexing, the mixture was oxygenated and incubated at 37 °C. After 4 h, the mixture was extracted using the modified Bligh and Dyer method^55^.Briefly, chloroform (1 mL) and methanol (1 mL) were then added. The sample was vortexed for 1 min and centrifuged at 500 × g for 5 min. The lower organic phase was collected, dried under a stream of nitrogen, and reconstituted in methanol containing 100 µM BHT.

Non-oxidized and oxidized SAPI samples were diluted to 100 µM in methanol and analyzed by LC– MS/MS as described below. The residual non-oxidized fraction was determined from the MS peak area of PI 38:4 and expressed relative to that of non-oxidized SAPI.

### PI4K2A activity assay

PI4K2A activity was measured using the ADP-Glo Kinase Assay (Promega, V6930). Recombinant active human PI4K2A protein (Abcam, ab268860; 10 ng per well) was incubated with non-oxidized or oxidized SAPI (0.2 µg per well) in 384-well plates for the indicated durations. The assay was performed according to the manufacturer’s instructions, and luminescence was measured using an EnSpire Multimode Plate Reader (PerkinElmer).

### PI4K2A S-palmitoylation assay

Calu-1 cells stably expressing TMEM192–3×HA were pretreated with 2-BP (20 µM) for 24 h, where indicated, and subsequently treated with LLOMe (1 mM) for 30 min or RSL3 (0.1 µM) for 3 h, followed by Lyso-IP. PI4K2A S-palmitoylation was analyzed using the RapidSPALM Protein S-Palmitoylation Detection Kit (BioDynamics Laboratory, F017A) according to the manufacturer’s instructions. Briefly, native S-palmitoyl groups were replaced with a multifunctional tag (MfTag), and MfTag-labelled proteins were enriched using a Loop affinity column. Enriched proteins were eluted using the supplied reducing reagent and analyzed by immunoblotting for PI4K2A. The proportion of S-palmitoylated PI4K2A was calculated by dividing the intensity of the palmitoylated band by the combined intensities of the palmitoylated and non-palmitoylated bands.

### PI4K2A phosphorylation assay

PI4K2A phosphorylation was analyzed by Phos-tag SDS–PAGE^56^. Equal amounts of protein were separated on polyacrylamide gels containing 20 µM Phos-tag Acrylamide (FUJIFILM Wako Pure Chemical Corporation, AAL-107S1). After electrophoresis, the gels were gently agitated three times for 10 min in transfer buffer containing 10 mM EDTA, followed by incubation for 10 min in EDTA-free transfer buffer. Proteins were transferred onto PVDF membranes under the same conditions used for conventional western blotting. Membranes were blocked with Blocking One-P (Nacalai Tesque) for 20 min and subjected to immunoblotting. The proportion of phosphorylated PI4K2A was calculated by dividing the intensity of the phosphorylated band by the combined intensities of the phosphorylated and non-phosphorylated bands.

### Western blotting

Cells were washed with PBS and lysed on ice for 30 min in buffer containing 50 mM Tris–HCl (pH 7.5), 150 mM NaCl, 1% Triton X-100, 0.1% SDS, 1% sodium deoxycholate, 1 mM benzylsulfonyl fluoride, 1 mM sodium orthovanadate, and protease inhibitor cocktail. Cell lysates were collected by scraping and centrifuged at 10,000 × g for 15 min. Protein concentrations were determined using a Protein Assay BCA Kit (Nacalai Tesque). Equal amounts of protein were mixed with 5× SDS sample buffer containing 50% glycerol, 0.05% bromophenol blue, 150 mM Tris–HCl (pH 6.8), 5% SDS, and 25% 2-mercaptoethanol. Proteins were separated by SDS–PAGE using 10% or 15% polyacrylamide gels and transferred onto PVDF membranes (Merck Millipore). Membranes were blocked with Blocking One (Nacalai Tesque) for 30 min at room temperature and incubated overnight at 4 °C with primary antibodies diluted in Blocking One. After three washes with TBS-T, the membranes were incubated with the appropriate secondary antibodies for 1 h at room temperature. Signals were developed using WSE-7120 EzWestLumi Plus (ATTO) and detected using a ChemiDoc MP system (Bio-Rad). Band intensities were quantified using Image Lab software (Bio-Rad). Mouse anti-PI4K2A antibody (Santa Cruz Biotechnology, sc-390026) was used at a dilution of 1:1,000.

### APAP-induced liver injury model

C57BL/6J mice were intraperitoneally administered APAP (300 mg/kg) dissolved in saline after overnight fasting, as described previously^21^. Blood was collected by cardiac puncture 4 h after APAP administration, after which the liver was excised. For Lip-1 treatment, Lip-1 was dissolved in 2.9% (v/v) DMSO in saline and administered intraperitoneally at 10 mg/kg 1 h after APAP administration. Ten minutes before sample collection, mice were anesthetized by intraperitoneal injection of a mixture of medetomidine hydrochloride (0.3 mg/kg), midazolam (4 mg/kg), and butorphanol tartrate (5 mg/kg) at a volume of 10 µL/g body weight. Liver tissues were immediately frozen in liquid nitrogen and stored at −80 °C until use. All animal experiments were conducted in accordance with the institutional guidelines for animal experimentation and were approved by the Committee on Ethics of Animal Experiments of Kyushu University (approval no. A25-157-2). Male C57BL/6J mice aged 7 weeks were purchased from Japan SLC. Mice were housed under a 12-h light/12-h dark cycle at 24 ± 1 °C and 60 ± 10% humidity, with free access to CLEA Rodent Diet CE-2 chow (CLEA Japan) and tap water. Before use, mice were acclimated for 1 week in the animal facility of the Faculty of Pharmaceutical Sciences, Kyushu University.

### Hepatocyte-specific conditional *Gpx4* knockout mouse

For generating tamoxifen-inducible hepatocyte-specific *Gpx4* knockout mice, *Alb-CreER^T^*^2^*;Gpx4^fl/fl^*mice (female, 11 months old) were used. *Alb-CreER^T^*^2^*;Gpx4^fl/fl^*mice were obtained by breeding *Gpx4^fl/fl^* mice with *Alb-CreER^T^*^2^ mice as previously described^24^. Cre-negative *Gpx4^fl/fl^ littermates* served as controls. To induce hepatocyte-specific deletion of *Gpx4*, the mice were intraperitoneally administered 2 mg tamoxifen (Nacalai Tesque, 19885-34) dissolved in triolein (Tokyo Chemical Industry Co., Ltd., G0089) on two consecutive days. At the time of the first tamoxifen injection, diet was changed from standard chow (Funabashi Farm Co., Ltd) to a vitamin E-deficient diet (<7 mg/kg vitamin E; Funabashi Farm Co., Ltd), as previously described^24, 57^. Mice were housed under controlled environmental conditions: temperature of 24 ± 3 °C, humidity of 50 ± 10%, and a 12 h light/dark cycle (lights on: 08:00–20:00, lights off: 20:00–08:00). Mice were euthanized one month after the first tamoxifen administration, and liver tissues were collected, immediately frozen in liquid nitrogen, and stored at −80 °C until analysis. All animal experiments were conducted in accordance with the Institutional Animal Care and Use Committees of Kyoto University (approved No. Med Kyo 26043-2), the Kyoto University Recombinant DNA Experiment Safety Committee (approved No. 260259), and the German Animal Welfare Law and have been approved by the institutional committee on animal experimentation and the government of Upper Bavaria (approved No. ROB-55.2-2532-Vet_02-18-13).

### Lipid extraction from tissues

Hepatic lipids were extracted from the liver samples according to the modified Bligh and Dyer method^55^. Briefly, methanol containing 100 μM BHT and 100 nM LysoPC 18:1(d7) was added to a frozen tissue sample (wet weight: 80 ± 10 mg), and then the sample was homogenized using a Macro Smash homogenizer. Subsequently, the extraction solutions were sonicated on an ice bath for 5 min.

After centrifugation (6000 × g, 10 min, 4 °C), 600 µL of the supernatant was collected and then 600 µL chloroform and 480 µL water were added to the supernatant. The organic layer was collected in a tube and dried under a stream of nitrogen gas; the dried residue was dissolved in 300 µL methanol and passed through a 0.45-µm Millex-LH filter (Merck Millipore). Samples were stored at −80 °C until analysis.

### Measurement of plasma ALT and AST activities

Plasma alanine aminotransferase (ALT) and aspartate aminotransferase (AST) activities were measured using the Transaminase CII-Test Wako kit (FUJIFILM Wako Pure Chemical Corporation, 431-30901) according to the manufacturer’s instructions.

### TBARS assay

Liver tissue (80 ± 10 mg) was homogenized by sonication on ice in 1 mL RIPA buffer containing 50 mM Tris–HCl (pH 7.5), 150 mM NaCl, 0.1% SDS, 1% Triton X-100, and 1% sodium deoxycholate. After incubation on ice for 5 min, homogenates were centrifuged at 16,000 × g for 5 min at 4 °C, and the supernatants were collected. A 100-µL aliquot of each supernatant was mixed with 10 µL BHT (50 mg/mL), 35 µL SDS (8%), 20 µL disodium EDTA (80 mM), 250 µL acetic acid (20%), and 250 µL thiobarbituric acid (0.8%; Nacalai Tesque, 33614-92). After mixing, samples were heated in a boiling-water bath for 60 min.

Samples were cooled on ice, mixed with 1 mL n-butanol/pyridine (15:1, v/v), and centrifuged at 800 × g for 10 min at 4 °C. A 200-µl aliquot of the upper phase was collected, and fluorescence was measured at excitation and emission wavelengths of 515 and 553 nm, respectively. Malondialdehyde levels were calculated using a calibration curve generated with 1,1,3,3-tetraethoxypropane.

### Histological analysis

Paraffin embedding of liver tissues was performed by the Department of Anatomic Pathology, Graduate School of Medical Sciences, Kyushu University. Paraffin blocks were sectioned at a thickness of 5 µm using a microtome (Leica). Sections were washed three times with xylene and rehydrated sequentially in 100% ethanol twice, 90% ethanol, and 70% ethanol. After washing with water for 2 min, sections were stained with hematoxylin solution (FUJIFILM Wako Pure Chemical Corporation, 131-09665) for 5 min, washed with water for 10 min, and stained with eosin solution (FUJIFILM Wako Pure Chemical Corporation, 050-06041) for 3 min. Sections were washed with purified water, dehydrated sequentially in 70% ethanol and 100% ethanol twice, and washed three times with xylene. After drying, sections were mounted using VectaMount Mounting Medium.

Images were acquired using a BZ-X810 microscope (Keyence).

### RT-qPCR

Total RNA was extracted from frozen liver tissues using the NucleoSpin RNA XS kit (MACHEREY-NAGEL). First-strand cDNA was synthesized from 500 ng of total RNA using ReverTra Ace qPCR RT Master Mix with gDNA Remover (TOYOBO, 11896-34). Real-time PCR was performed using the THUNDERBIRD Next SYBR qPCR Mix (TOYOBO, 20498-74) and the CFX Duet Real-Time PCR System (Bio-Rad) according to the manufacturer*’*s instructions. The primer sequences were as follows: Actb, forward 5’-GGCTGTATTCCCCTCCATCG-3’ and reverse 5’-CCAGTTGGTAACAATGCCATGT-3’; Gpx4, forward 5’-TTACGAATCCTGGCCTTCCC-3’ and reverse 5’-CCACGCAGCCGTTCTTATCA-3’.

### LC–MS/MS analysis of lipid peroxidation products

Lipid extracts from cultured cells, tissues, and lysosomal fractions were analyzed using an LCMS-8060 triple-quadrupole mass spectrometer (Shimadzu) equipped with an electrospray ionization source. Chromatographic separation was performed under the following conditions, which were also used for LC–HRMS and HRMS/MS analyses: injection volume, 5 µL; autosampler temperature, 4 °C; column, InertSustain C18 (2.1 × 150 mm, 3-µm particle size; GL Sciences); column temperature, 40 °C; mobile phase A, 5 mM ammonium formate in acetonitrile/water (2:1, v/v); mobile phase B, 5 mM ammonium formate in isopropanol/methanol (19:1, v/v); flow rate, 0.4 mL/min; and gradient, 0–22.5 min, 0–100% B, and 22.5–27.5 min, 100% B.

MS analysis was performed in positive-ion or negative-ion mode using multiple-reaction monitoring transitions optimized for the individual LPO products. The source settings were as follows: nebulizing gas flow, 3 L/min; heating gas flow, 10 L/min; interface temperature, 300 °C; desolvation line temperature, 250 °C; heat-block temperature, 400 °C; and drying gas flow, 10 L/min. MS peak areas were normalized to LysoPC 18:1(d7) as an internal standard where indicated. Data were acquired using LabSolutions 5.135 (Shimadzu) and processed using Multi-ChromatoAnalysT 1.4.0.0 software (Beforce).

### LC–HRMS and HRMS/MS analyses

LC–HRMS and HRMS/MS analyses were performed using a quadrupole Orbitrap mass spectrometer (Thermo Fisher Scientific) under the chromatographic conditions described above. For positive-ion mode, the source settings were as follows: sheath gas flow, 40 arbitrary units; auxiliary gas flow, 10 arbitrary units; spray voltage, 3,500 V; capillary temperature, 250 °C; S-lens RF level, 50; and heater temperature, 425 °C. For negative-ion mode, the source settings were identical except that the spray voltage was 2,000 V.

Full-scan HRMS data were acquired at a resolving power of 70,000, with an automatic gain control target of 3 × 10^6^, a maximum injection time of 200 ms and a scan range of *m/z* 200–1,500.

HRMS/MS data were acquired at a resolving power of 17,500, with an automatic gain control target of 3 × 10^6^, a maximum injection time of 200 ms, an isolation window of ±0.4 Da, a fixed first mass of *m/z* 50 and a normalized collision energy of 30.

### Structural characterization of PI 18:0_20:4;O3

Targeted LC–MS/MS analysis of PI 18:0_20:4;O3 was performed using an Orbitrap Exploris 240 mass spectrometer equipped with a heated electrospray ionization source and controlled by Xcalibur software (Thermo Fisher Scientific). Chromatographic separation was performed using a Vanquish/Dionex Ultimate 3000 LC system. Separation was achieved on an InertSustain C18 (2.1 × 150 mm, 3-µm particle size). The flow rate was set to 0.400 mL/min. The LC gradient was programmed as follows: 0–22.5 min, 0–100% solvent B; 22.5–27.5 min, 100% solvent B; 27.5–27.6 min, 100–0% solvent B; and 27.6–30.0 min, 0% solvent B for re-equilibration.

The mass spectrometer was operated in negative ion mode using a targeted SIM-dependent MS/MS method. The sheath gas, auxiliary gas, and sweep gas flow rates were set to 40, 10, and 1 arbitrary units, respectively. The ion transfer tube and vaporizer temperatures were maintained at 275°C and 300°C, respectively. Targeted SIM acquisition was performed from 1.6 to 30 min with an isolation window of 1 *m/z*. The Orbitrap resolution for the targeted SIM scan was set to 60,000, with an RF lens value of 70%, a standard AGC target, automatic maximum injection time, one microscan, and profile data acquisition. The target precursor ion for PI 18:0_20:4;O3 was monitored as the deprotonated ion [M–H]⁻ at *m/z* 933.5346, corresponding to the formula C47H83O16P.

Data-dependent MS/MS spectra were acquired with up to 20 dependent scans. Precursor ions were isolated with a 2 *m/z* isolation window and fragmented by higher-energy collisional dissociation using a normalized collision energy of 30%. MS/MS spectra were acquired at an Orbitrap resolution of 15,000 over an *m/z* range of 50–950. The AGC target was set to standard, the maximum injection time was set to automatic, and data were acquired in profile mode. Oxidized lipid data were analyzed using Qual Browser software (Thermo Fisher Scientific, San Jose, CA, USA).

### LC–MS/MS analysis of PI(4)P

The phosphoinositide analysis was conducted at Lipidome Lab Co., Ltd, as described previously^58^. Briefly, samples were dissolved in methanol, and total lipids were extracted by the modified Bligh and Dyer method. The total lipid fraction was further purified and recovered using an anion exchange column to isolate the acidic phospholipid fraction, including phosphoinositides. The purified fraction was dried under a stream of nitrogen, then redissolved in acetonitrile and transferred to vials. LC-MS/MS analysis was performed using the Xevo TQ-XS mass spectrometer with an ACQUITY UPLC H-Class (Waters). The lipids were separated using a CHIRALPAK IC-3 column (2.1 × 250 mm, 3 mm, DAICEL) at 23°C under the conditions as described^58^.

Phosphoinositide species were measured using MRM in positive ion mode, and peak picking was conducted using analytical software MassLynx4.2 (Waters). Peak areas of individual species were normalized with those of the internal/surrogate standards, and final quantification data were normalized to the total cell number and expressed as pmol per 1 × 10^6^ cells.

### Statistical analysis

Data are presented as the mean ± s.d. unless otherwise indicated. Statistical significance was assessed using a two-sided unpaired t-test, Mann–Whitney U test, one-way analysis of variance followed by Dunnett’s test or the Tukey–Kramer test, two-way analysis of variance followed by Sidak’s multiple-comparisons test, or the Kruskal–Wallis test, as indicated in the figure legends. Statistical analyses were performed using GraphPad Prism version 9.5.1 (GraphPad Software).

## ACKNOWLEDGMENTS

This study was partly supported by JSPS KAKENHI (23H05481, 24K22024, 25K24600, and 26K23324 to KY) grants: the Takeda Science Foundation (to KY) : the Ono Medical Research Foundation (to KY) : the Nagase Science and Technology Foundation (to KY) : the Hoansha Foundation (to KY). M.C. received funding from the European Research Council (ERC) under the European Union’s Horizon 2020 research and innovation program (grant agreement no. GA 884754). The funding sources had no role in the study design, data collection and analysis, decision to publish, or preparation of the manuscript. We appreciate the technical assistance from the Research Support Center, Research Center for Human Disease Modeling, and the Autonomous Medical Research Center, Kyushu University Graduate School of Medical Sciences. The Research Support

Center is partially supported by the Mitsuaki Shiraishi Fund for Basic Medical Research.

## AUTHOR CONTRIBUTIONS

S.H., P.Y.H., Y.J., H.N., K.M., Y.S., M.Y., H.N., and performed and analyzed the cell experiments. Y.J., Y.M., E.K., T.U., and N.K. performed and analyzed the animal experiments. S.T., and H.M. synthesized the oxPI. S.H., P.Y.H., Y.J., Y.M., E.K., Y.S., M.J., M.A., P.L., M.C., and G.H. contributed scientific insights and analyzed the results. S.H., P.Y.H., and Y.M. wrote parts of the manuscript. K.Y. designed the experiments, conceived and supervised the study, and wrote and edited the manuscript. All authors have read and agreed to the contents of the paper.

## FIGURE LEGENDS

**Supplementary Fig. 1.**
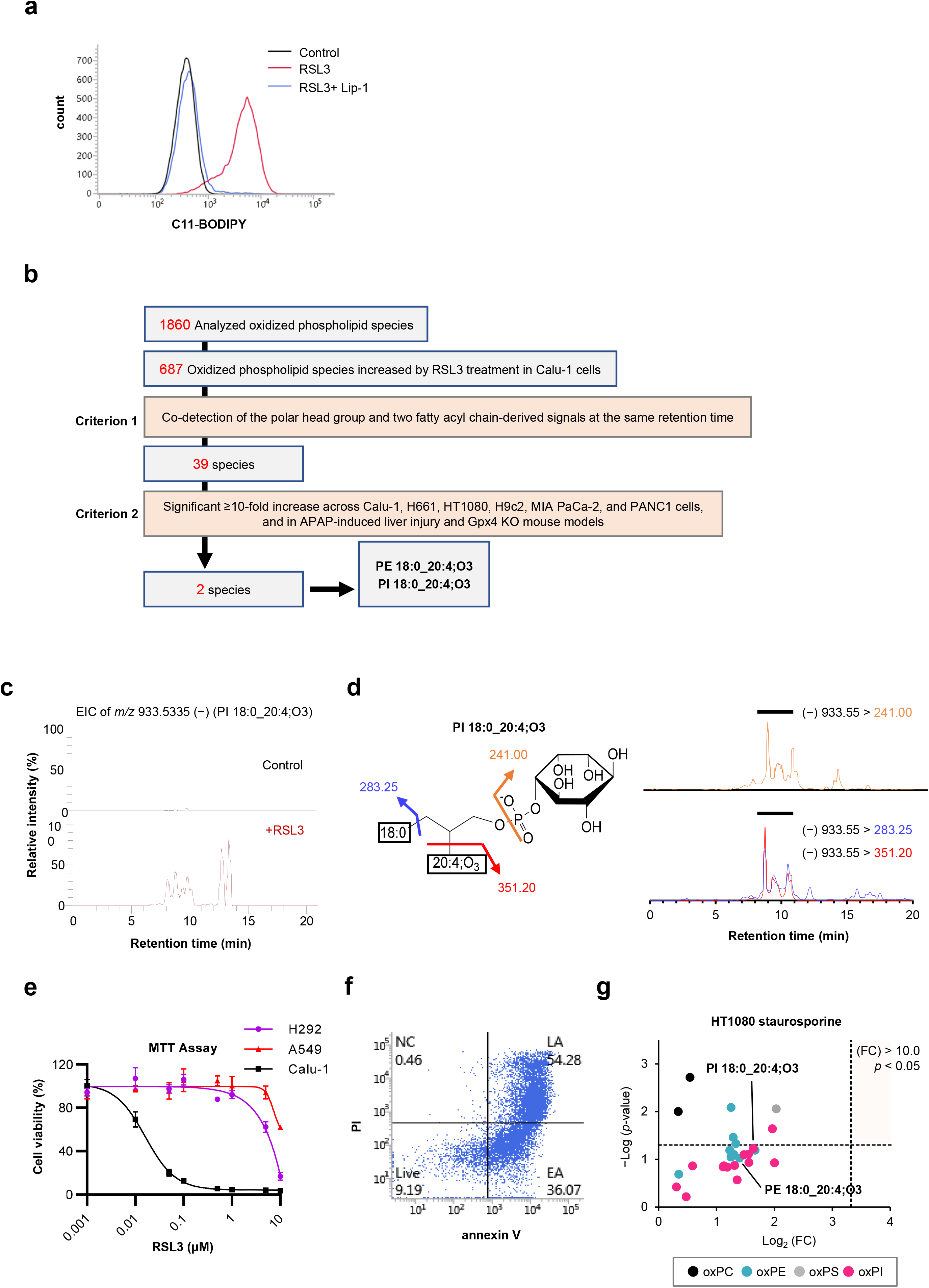
Comprehensive analysis of oxidized phospholipids generated during ferroptosis. **a,** Representative flow cytometry plot showing LPO detected using C11-BODIPY. Calu-1 cells were treated with RSL3 (0.1 µM) with or without Lip-1 (1 µM) for 5 h. **b,** Workflow for screening and identifying oxidized phospholipid species commonly increased across six cell lines, APAP-induced liver injury model mice, and *Alb-CreER^T^*^2^*;Gpx4^fl/fl^* mice. **c,** Representative EICs at *m/z* 933.5335 obtained by LC–HRMS in Calu-1 cells treated with RSL3 (0.1 µM) for 5 h. **d,** LC–MS/MS analysis of three product ions derived from oxidized phospholipids. Co-eluted peaks were detected at retention times of 8–11 min, and the lipid species was assigned as PI 18:0_20:4;O3. **e,** Viability of Calu-1, H292 and A549 cells treated with RSL3 (0.0001–10 µM) for 24 h, as determined using the MTT assay. **f,** Representative flow cytometry plot of annexin V–FITC and propidium iodide fluorescence in STS-treated HT1080 cells. Apoptosis was induced by treatment with STS (0.1 µM) for 24 h, followed by STS (1 µM) for 4 h. **g,** Volcano plot showing changes in oxidized phospholipid abundance in STS-treated HT1080 cells relative to control cells. Data in (**e)** are presented as the mean ± s.d. of three independent experiments. Data in (**g)** were obtained from three independent experiments, and statistical significance was assessed using a two-sided unpaired t-test.

**Supplementary Fig. 2.**
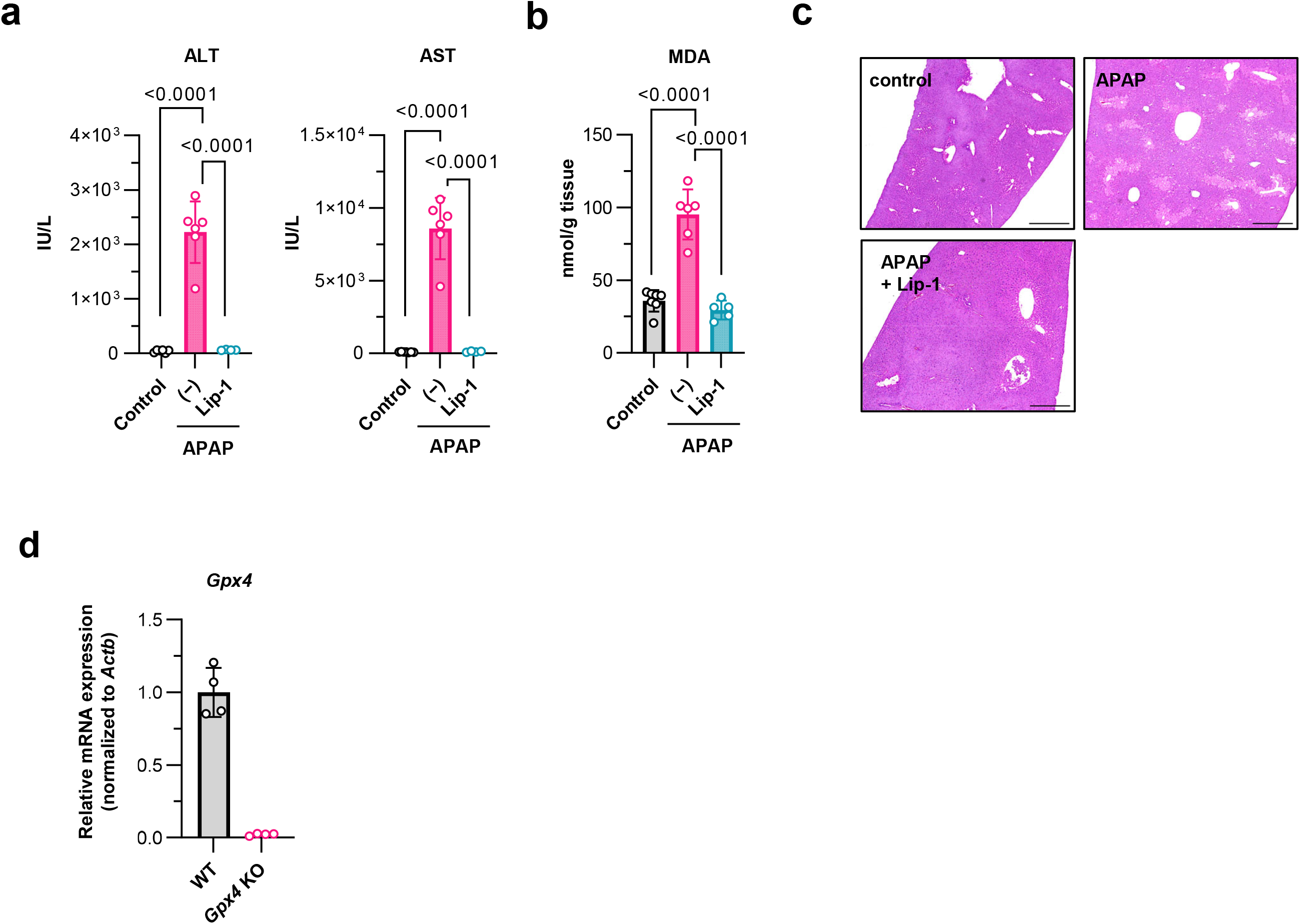
Validation of acetaminophen (APAP)-induced liver injury and hepatocyte-specific *Gpx4* conditional knockout model. **a–c,** Mice were administered APAP (300 mg/kg, intraperitoneally), followed by Lip-1 (10 mg/kg, intraperitoneally) 1 h later. Plasma and liver tissues were collected 4 h after APAP administration. **a,** Plasma ALT and AST activities. **b,** Hepatic MDA levels measured using the TBARS assay. **c,** Representative H&E-stained liver sections. **d,** Relative *Gpx4* mRNA expression in liver tissues from WT and *Alb-CreER^T^*^2^*;Gpx4^fl/fl^* mice (n = 4 mice per group). Data in (**a, b**) are mean ± s.d.; n = 4–7 mice per group, by the Tukey–Kramer test. Scale bars, 500 µm (**c**).

**Supplementary Fig. 3.**
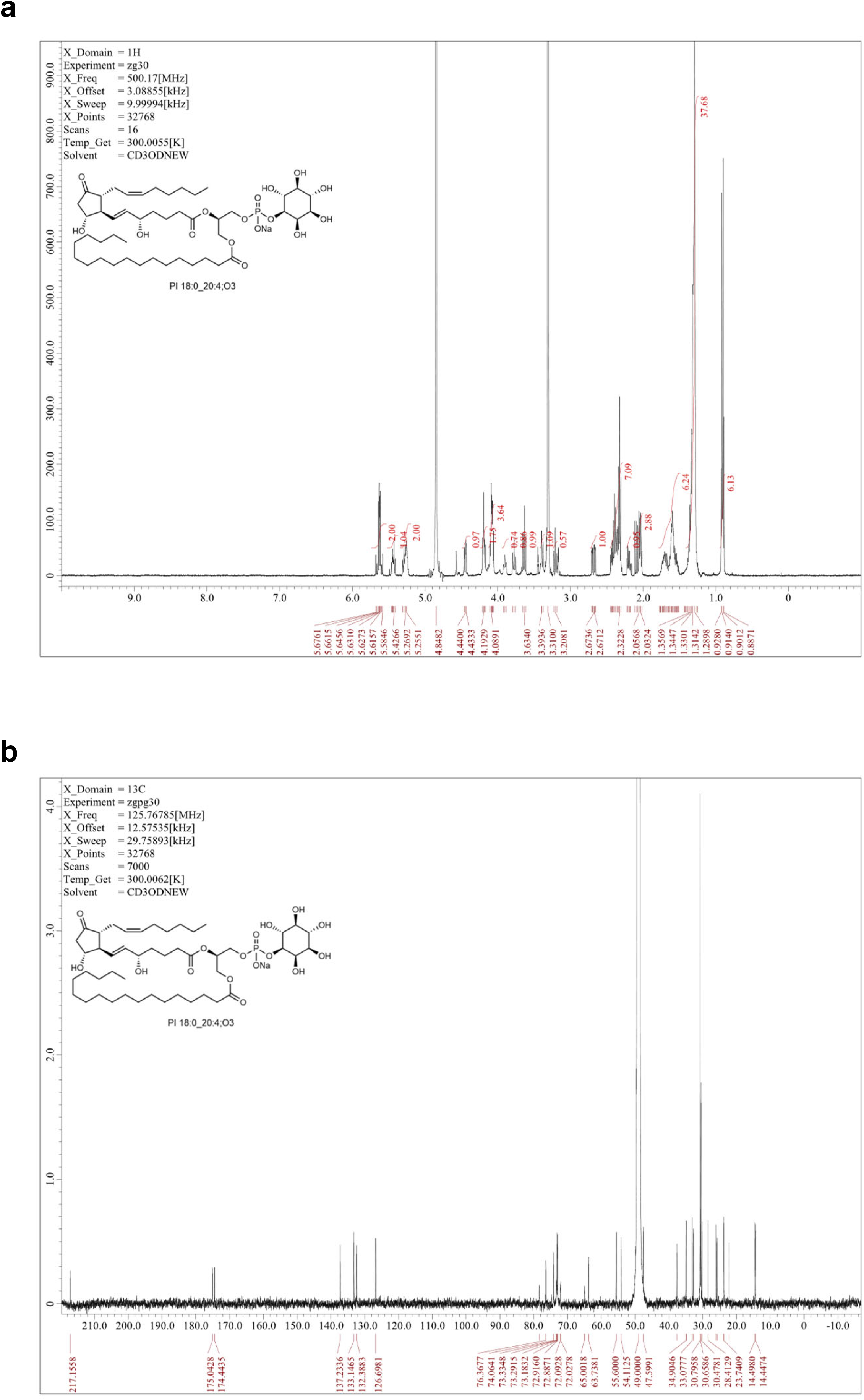
Synthesis of PI 18:0/20:1(13E);[8–12cy5;11OH;9oxo];15OH (PI 18:0_20:4;O3). **a,** ^1^H NMR (500 MHz, CD_3_OD): δ 5.65 (dd, *J* = 15.3, 7.3 Hz, 1H), 5.61 (dd, *J* = 15.3, 6.4 Hz, 1H), 5.47–5.40 (m, 1H), 5.32–5.22 (m, 2H), 4.45 (dd, *J* = 11.9, 3.4 Hz, 1H), 4.23–4.15 (m, 2H), 4.12– 4.04 (m, 4H), 3.90 (ddd, *J_H-H_* = 9.0, 9.0, *J_H-P_* = 2.8 Hz, 1H), 3.77 (dd, *J* = 9.5, 9.5 Hz, 1H), 3.63 (dd, *J* = 9.7, 9.5 Hz, 1H), 3.39 (dd, *J* = 9.7, 2.4 Hz, 1H), 3.21 (dd, *J* = 9.5, 9.5 Hz, 1H), 2.68 (ddd, *J* = 18.6, 7.3, 1.2 Hz, 1H), 2.46–2.29 (m, 7H), 2.20 (ddd, *J* = 11.3, 5.8, 5.5 Hz, 1H), 2.08 (dd, *J* = 18.6, 9.5 Hz, 1H), 2.04 (td, *J* = 7.3, 6.7 Hz, 2H), 1.79–1.49 (m, 4H), 1.40–1.23 (m, 34H), 0.91 (t, *J* = 7.0 Hz, 3H), 0.90 (t, *J* = 7.0 Hz, 3H). HRMS(ESI): *m/z* [M+Na]^+^ calcd for C_47_H_82_Na_2_O_16_P 979.5136, found 979.5146. [α]_D_^24^ -3.8 (c = 0.15, CHCl_3_). **b,** ^13^C{^1^H} NMR (126 MHz, CD_3_OD): δ 217.16, 175.04, 174.44, 137.23, 133.15, 132.39, 126.70, 78.40, 76.37, 74.06, 73.31 (d, *J_C-P_* = 5.5 Hz), 73.18 (2C), 72.92, 72.89, 72.06 (d, *J_C-P_* = 8.2 Hz), 64.98 (d, *J_C-P_* = 4.5 Hz), 63.74, 55.60, 54.11, 47.60, 37.67, 34.93, 34.90, 33.08, 32.76, 30.80 (8C), 30.66, 30.48 (3C), 30.23, 28.41, 26.01, 25.78, 23.74, 23.69, 22.15, 14.50, 14.45. HRMS(ESI): *m/z* [M+Na]^+^ calcd for C_47_H_82_Na_2_O_16_P 979.5136, found 979.5146. [α]_D_^24^ -3.8 (c = 0.15, CHCl_3_).

**Supplementary Fig. 4.**
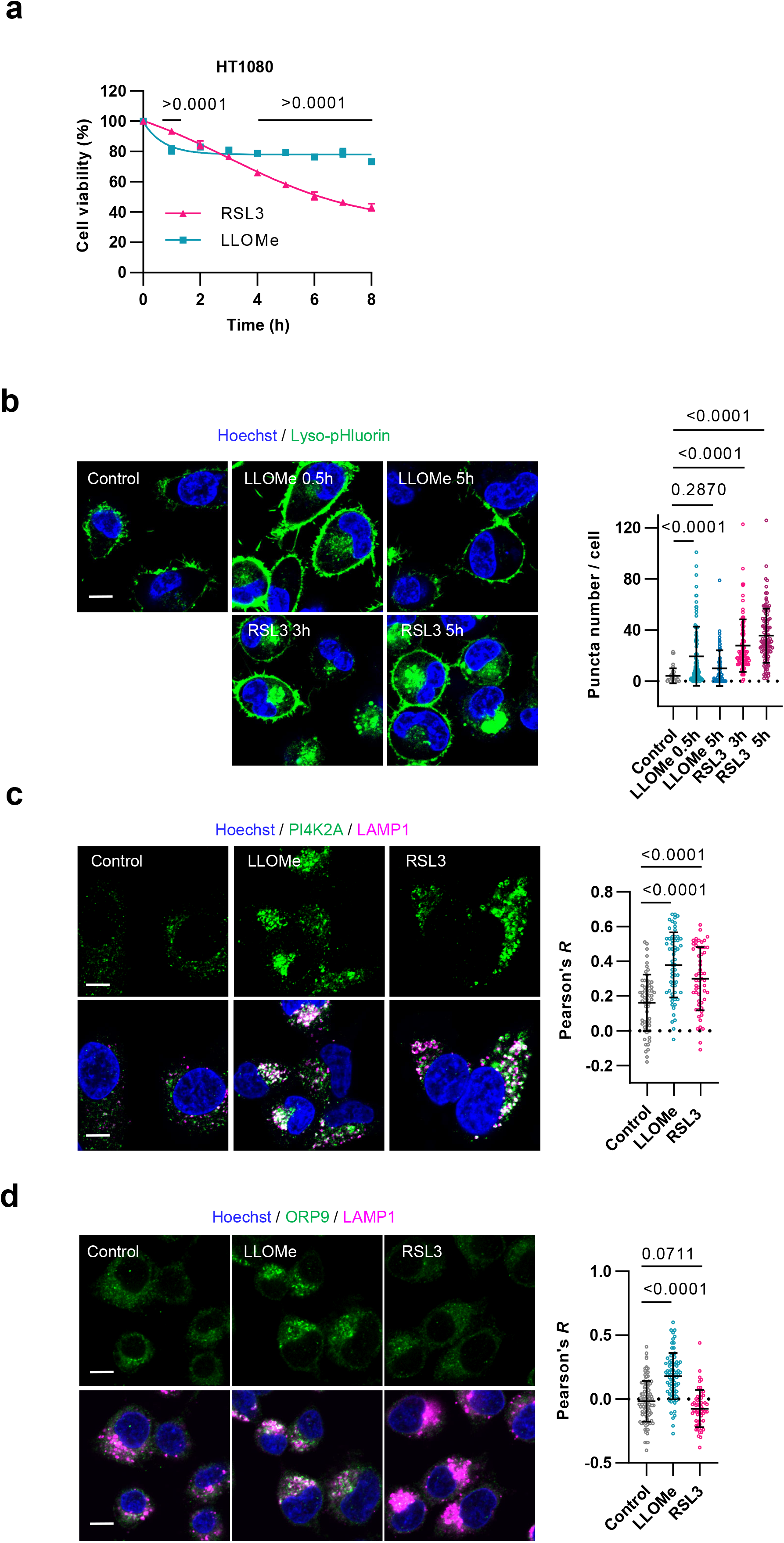
Ferroptosis impairs PITT-mediated lysosomal membrane repair in HT1080 cells. **a,** Viability of HT1080 cells treated with LLOMe (1 mM) or RSL3 (0.1 µM) for the indicated durations (1–8 h). **b,** HT1080 cells stably expressing lyso-pHluorin were treated with LLOMe (1 mM) or RSL3 (0.1 µM) for the indicated durations. The number of lyso-pHluorin puncta per cell was quantified. **c,** HT1080 cells were treated with LLOMe (1 mM) for 30 min or RSL3 (0.1 µM) for 3 h and immunostained for PI4K2A and LAMP1. **d,** HT1080 cells were treated with LLOMe (1 mM) for 30 min or RSL3 (0.1 µM) for 3 h and immunostained for ORP9 and LAMP1. Colocalization with LAMP1 was quantified using Pearson’s correlation coefficient (**c–d**). Data in (**a)** are presented as the mean ± s.d. of three independent experiments. Statistical significance was assessed using Sidak’s multiple-comparisons test. Data in (**b-d)** are presented as the mean ± s.d. Each data point represents an individual cell quantified from three independent images; n = 44–101 (**b**), 55–60 (**c**), 53–109 (**d**) cells. Statistical significance was assessed using Dunnett’s test. Scale bars, 10 µm (**b–d**).

**Supplementary Fig. 5.**
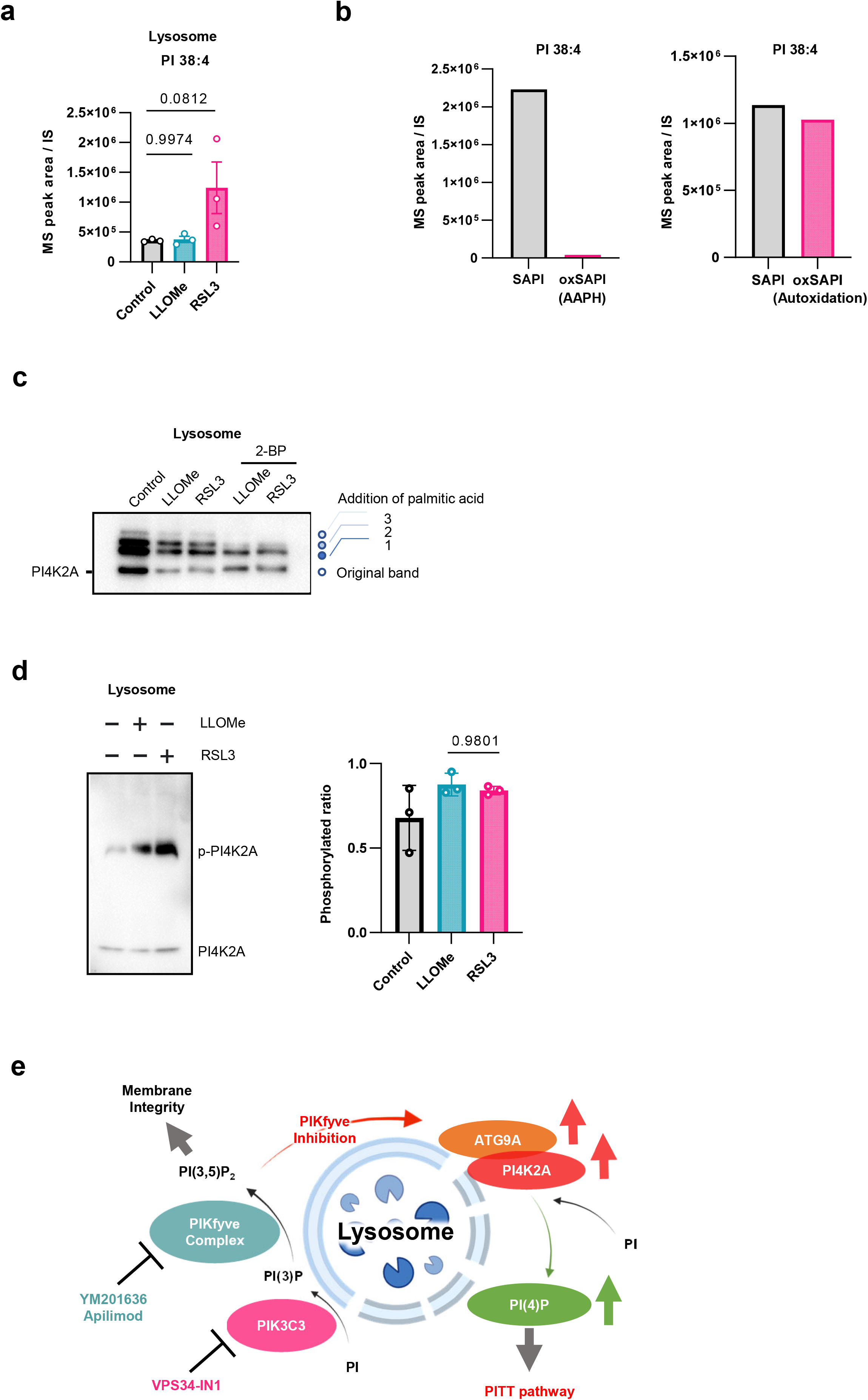
Additional analyses of PI4K2A activity and PI(4)P-dependent lysosomal membrane repair. **a,** LC–MS/MS-based lysosomal lipidomic analysis. Calu-1 cells stably expressing TMEM192– 3×HA were treated with LLOMe (1 mM, 30 min) or RSL3 (0.1 µM, 3 h), followed by Lyso-IP. PI 38:4 in lysosomal fractions were quantified and normalized to the internal standard. n = 3. **b,** LC– MS/MS analysis of residual non-oxidized PI 38:4 in AAPH-oxidized SAPI and autoxidized SAPI relative to non-oxidized SAPI. Residual PI 38:4 was quantified from MS peak areas. n = 1. **c,** PI4K2A S-palmitoylation assay. Calu-1 cells stably expressing TMEM192–3×HA were pretreated with 2-BP (20 µM) for 24 h and subsequently treated with LLOMe (1 mM) for 30 min or RSL3 (0.1 µM) for 3 h, followed by Lyso-IP. Lysosomal fractions were analyzed using the RapidSPALM Protein S-Palmitoylation Detection Kit, and PI4K2A S-palmitoylation was assessed by immunoblotting. The proportion of S-palmitoylated PI4K2A was calculated by dividing the intensity of the palmitoylated band by the combined intensities of the palmitoylated and non-palmitoylated bands. n = 2 for the control, LLOMe and RSL3 groups; n = 1 for the LLOMe plus 2-BP and RSL3 plus 2-BP groups. **d,** PI4K2A phosphorylation assay. Calu-1 cells stably expressing TMEM192– 3×HA were treated with LLOMe (1 mM) for 30 min or RSL3 (0.1 µM) for 3 h, followed by Lyso-IP. PI4K2A phosphorylation was assessed by Phos-tag immunoblotting. The proportion of phosphorylated PI4K2A was calculated by dividing the intensity of the phosphorylated band by the combined intensities of the phosphorylated and non-phosphorylated bands. Data in (**d)** are presented as the mean ± s.d. of three independent experiments. Statistical significance was assessed using the Tukey–Kramer test. **e,** Schematic illustration of PIKfyve inhibition. Created in BioRender. Yamada, K. (2027) https://BioRender.com/z5nmsby.

